# Therapeutic modulation of the blood-brain barrier and ischemic stroke by a bioengineered FZD_4_-selective WNT surrogate

**DOI:** 10.1101/2022.10.13.510564

**Authors:** Jie Ding, Sung-Jin Lee, Lukas Vlahos, Kanako Yuki, Cara C. Rada, Vincent van Unen, Meghah Vuppalapaty, Hui Chen, Asmiti Sura, Aaron K. McCormick, Madeline Tomaske, Samira Alwahabi, Huy Nguyen, William Nowatzke, Lily Kim, Lisa Kelly, Douglas Vollrath, Andrea J. Califano, Wen-Chen Yeh, Yang Li, Calvin J. Kuo

## Abstract

Derangements of the blood-brain barrier (BBB) or blood-retinal barrier (BRB) occur in disorders ranging from stroke, cancer, diabetic retinopathy, and Alzheimer’s disease. The Norrin/FZD_4_/TSPAN12 pathway activates WNT/β-catenin signaling, which is essential for BBB and BRB function. However, systemic pharmacologic FZD_4_ stimulation is hindered by obligate palmitoylation and insolubility of native WNTs and suboptimal properties of the FZD_4_-selective ligand Norrin. Here, we developed L6-F4-2, a non-lipidated, FZD_4_-specific surrogate with significantly improved sub-picomolar affinity versus native Norrin. In Norrin knockout (*Ndp^KO^*) mice, L6-F4-2 not only potently reversed neonatal retinal angiogenesis deficits, but also restored BRB and BBB function. In adult C57Bl/6J mice, post-stroke systemic delivery of L6-F4-2 strongly reduced BBB permeability, infarction, and edema, while improving neurologic score and capillary pericyte coverage. Our findings reveal systemic efficacy of a bioengineered FZD_4_-selective WNT surrogate during ischemic BBB dysfunction, with general applicability to adult CNS disorders characterized by an aberrant blood-brain barrier.

---

The cerebrovasculature comprises a highly specialized vascular bed that tightly regulates the movement of nutrients, ions, and cells from the blood into the brain parenchyma. This stringent cerebrovascular integrity is controlled by the blood-brain barrier (BBB) and blood-retinal barrier (BRB) to meet high metabolic demands, while specifically inhibiting transmigration of toxins and pathogens to prevent neuronal injury^1^. The BBB and BRB are comprised of a neurovascular unit (NVU) consisting of an endothelial cell (EC) layer with abundant tight junctions and covered by a dense pericyte layer, all engulfed by astrocyte end-feet^2^. BBB disruption and/or dysregulation is particularly significant to the pathology of numerous central nervous system (CNS) diseases including stroke, cancer, epilepsy, multiple sclerosis, ALS, Alzheimer’s disease, and recently, SARS-CoV-2 “brain fog”^3–8^.

Significant recent evidence has demonstrated essential WNT/β-catenin signaling regulation of both BBB/BRB function and CNS angiogenesis. The 19 ligands of the WNT family functionally crosslink Frizzled (FZD) 7-pass transmembrane/GPCR-like receptors to LRP5/6 co-receptors to initiate β-catenin signaling. Subsequently, inhibition of Axin-dependent degradation enables β-catenin nuclear translocation, association with LEF/TCF transcription factors and downstream target gene transactivation^9^. Within CNS endothelium, multiple inputs converge onto WNT/β-catenin signaling, including WNT binding to receptors as in the Norrin/FZD_4_/TSPAN12 and WNT7/GPR124/RECK pathways^10^. Mouse and zebrafish mutations in the aforementioned signaling nodes inhibit endothelial WNT/β-catenin signaling, with overlapping embryonic lethal phenotypes of impaired brain angiogenesis, glomeruloid vascular malformations, hemorrhage and BBB/BRB immaturity and leakage^11–14^. Importantly, compromise of endothelial or generalized WNT/β-catenin signaling exacerbates disease pathology in adult mice with hemorrhagic transformation of experimental stroke and glioma, with reversal by genetic WNT/β-catenin signaling activation^15–17^.

Frizzled-4 (FZD_4_) is particularly relevant for BBB/BRB regulation. FZD_4_ is highly expressed in CNS endothelium, while deletion in retinal or cerebellar EC elicits BRB compromise and retinal angiogenesis deficits in neonates and cerebellar BBB leakage in juvenile mice^10,11,18^. The secreted protein Norrin, encoded by *Ndp*, exhibits remarkable binding specificity for FZD_4_, but not other FZD family members. Despite lack of homology to classical WNTs, Norrin potently triggers FZD_4_ ubiquitination and internalization^19^, and activates β-catenin-dependent WNT/β-catenin signaling^20,21^ through coreceptors LRP5/6 and TSPAN12, while ectopic transgenic Norrin production restores cerebellar BBB integrity in *Ndp^KO^* mice^18^. Mutation of *Ndp*^22,23^ or its receptors FZD_4_^21,24^, *Lrp5*^25,26^ or *Tspan12*^27,28^, compromises EC WNT/β-catenin signaling, with neonatal ocular manifestations of impaired retinal angiogenesis, persistent hyaloid vessels, BRB leakage, and cerebellar BBB permeability. These mutations also underlie human vitreoretinal diseases, including Norrie disease, familial exudative vitreoretinopathy, and Coats’ disease^29,30^.

Despite significant potential, successful pharmacologic FZD_4_ activation for therapeutic BBB modulation has remained elusive. Norrin is a FZD_4_-selective surrogate but is a short-range signal that 3 is poorly secreted and highly associated with the extracellular matrix^31,32^. Unlike WNTs, which robustly activate WNT/β-catenin signaling with FZDs in the absence of co-transfected *LRPs*, Norrin requires both FZD_4_ and LRP5/6, implying that Norrin interacts in a ternary complex with FZD_4_ and LRPs^31,32^. WNT proteins themselves are obligately palmitoylated, which restricts expression, solubility, and in vivo pharmacokinetics^33,34^. In contrast, we developed bioengineered lipid-free FZD-specific WNT surrogates that signal by crosslinking FZD and LRP5/6 via anti-FZD single-chain antibodies (scFv) fused to LRP5/6-binding scFv or DKK1 C-terminal domains^35,36^. We further designed an array of FZD-selective surrogates active in a narrow dosing window, including monoselective FZD_4_ ligands that regulated hepatic zonation in vivo^37,38^. Chidiac et al. also described a FZD_4_-selective WNT surrogate (F4L5.13) that regulates BRB function in vivo with restoration of neonatal retinal barrier and angiogenesis defects in *Tspan12*^-/-^ mice, and partial rescue at 2-3 months of age^28^.

Here, we describe an improved FZD_4_-selective WNT surrogate (L6-F4-2) with subpicomolar affinity representing multi-log affinity improvements over Norrin and previously reported FZD_4_ surrogates^28,37^. L6-F4-2 bioactivity was confirmed in cultured brain EC and by in vivo rescue of neonatal retinal BRB function and angiogenesis defects in *Ndp^KO^* mice. Notably, L6-F4-2 also successfully treated blood-brain barrier pathology in mature mice by rescue of *Ndp^KO^* cerebellar BBB leakage. Crucially, we extended these findings to adult blood-brain barrier function using ischemic stroke as a model of prevalent disease, where post-stroke administration of L6-F4-2 reversed BBB permeability, infarction, edema, and neurologic score. Overall, these studies develop a highly optimized FZD_4_-selective WNT surrogate and validate FZD_4_ stimulation as a general therapeutic strategy for pathologic BBB compromise in neurological diseases.

## Results

### Generation of a sub-picomolar affinity monospecific FZD_4_-selective WNT surrogate, L6-F4-2

Given the genetic evidence supporting the importance of FZD_4_ in retinal vascular biology and BBB/BRB function, we generated a FZD_4_/LRP6-specific WNT mimetic molecule to crosslink these targets and initiate WNT/β-catenin signaling. The FZD_4_ binder selected was 5063, hereafter F4-2 (WO 2019/159084 A1). The fragment antigen-binding (Fab) form of F4-2 binding to the cysteine-rich domain (CRD) of FZD_4_ was confirmed by biolayer interferometry. The dissociation constant (K_D_) of the monovalent binding was observed as ~42 nM (Fig. 1a). As we and others previously showed that tetravalent bi-specific antibodies with two FZD and two LRP binding arms are highly potent and efficacious WNT mimetics in inducing β-catenin signaling^39,40^, we assembled F4-2 with the LRP6 binder, YW211.31.57^40^ (hereafter L6), in the tetravalent antibody format. This is referred to as L6-F4-2, where L6 is fused to the N-terminus of F4-2 IgG (Fig. 1b). Due to the avidity effect from the bivalent binding, L6-F4-2 showed extremely high affinity binding toward FZD_4_-CRD with K_D_ < 1 pM (Fig. 1b), which strongly surpasses previously described affinity measurements of native Norrin or previously described FZD_4_ surrogates (11-260 nM)^41,42^. Notably, L6-F4-2 FZD_4_ specificity was also preserved in the tetravalent format (Fig. 1c)

**Fig. 1.**
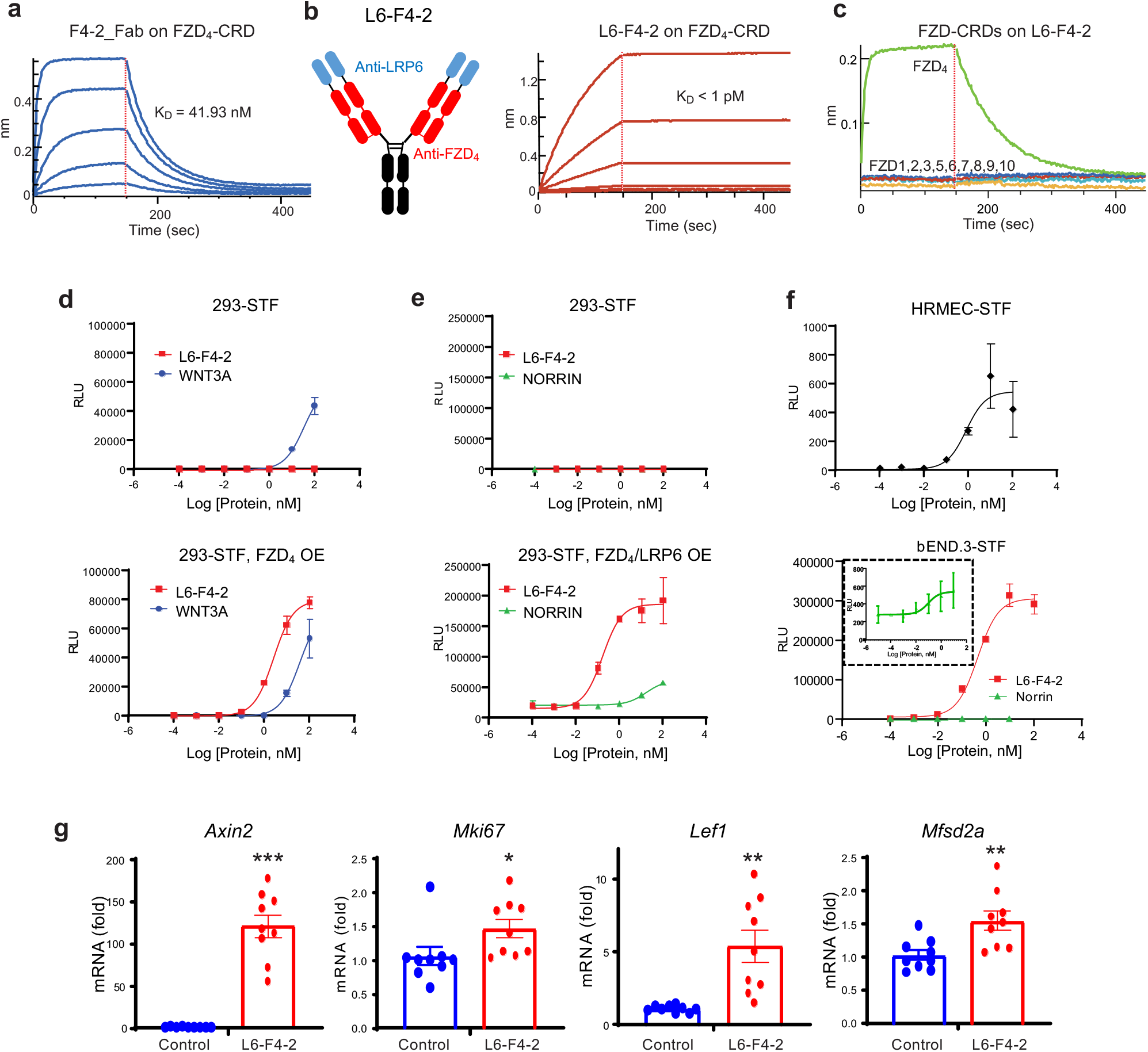
Characterization of the monoFZD_4_-specific WNT/β-catenin signaling surrogate, L6-F4-2. **a** Binding affinity of the recombinant F4-2_Fab molecule to FZD_4_ CRD measured by BLI assay. Dotted lines indicate the global fits generated by using a 1:1 Langmuir binding model. **b** Schematic of L6-F4-2 and the binding affinity of the L6-F4-2 molecule to the FZD_4_ CRD measured by BLI assay. Dotted lines indicate the global fits generated by using a 1:1 Langmuir binding model. **c** The binding specificity of L6-F4-2 against all ten FZD CRDs was examined by BLI assay. **d** Dose-dependent STF activity of L6-F4-2 or WNT3A in 293STF cells (top) or FZD_4_-transfected 293STF cells (bottom, OE=overexpression). **e** Dose-dependent STF activity of the L6-F4-2 or Norrin in 293STF cells (top) or cells transfected with both FZD_4_ and LRP6 (bottom, OE=overexpression). **f** Dose-dependent STF activities of L6-F4-2 in HRMEC cells (top) and bEnd.3 cells (bottom). Inset box in the bEnd.3-STF graph is an enlarged plot of the Norrin response. **g** Quantitative RT-PCR results of *Axin2*, *Ki67*, *Lef1* and *Mfsd2a* gene expression in bEnd.3 cells. mRNA expression values were normalized by *Acta2* gene expression. Results are from three independent experiments. Graphs are shown as mean ± SEM; *p < 0.05; ** p < 0.01; ***p < 0.001 Mann-Whitney U test.

The ability of L6-F4-2 to activate WNT/β-catenin signaling was assessed in WNT-responsive HEK293 Super TOP-FLASH (STF) reporter cells^36^ (293STF assay). While L6-F4-2 did not induce a strong response in parental 293STF cells due to low levels of endogenous FZD_4_, L6-F4-2 strongly and dose-dependently activated WNT/β-catenin signaling in FZD_4_-transfected 293STF cells (Fig. 1d). This activity was stronger than recombinant WNT3A (Fig. 1d) and recombinant Norrin in 293STF cells with FZD_4_ and LRP6 overexpression (Fig. 1e). A distinct monospecific FZD_4_-selective WNT surrogate, F4L5.13, was reported using an alternative molecular format, with dumbbell-like structures against two FZD_4_ and one or two LRP5 molecules and exhibited comparable maximal STF response to recombinant Norrin^28^. However, our monospecific FZD_4_ activator, L6-F4-2, was ~100 times more potent (EC50, L6-F4-2 0.1658 nM vs. Norrin 16.35 nM) with ~30 times higher E_max_ than Norrin (Fig. 1e). Though these different WNT mimetics were not directly compared side-by-side nor against the same Norrin preparation, these results suggest that affinity, appropriate molecular format, and geometry are critical for WNT mimetic generation^40^. We further performed STF assays in human retinal microvascular endothelial cells (HRMEC) and murine brain microvascular endothelial cells (bEND.3). As both of these EC lines express FZD_4_, L6-F4-2 induced dose-dependent STF responses in each without requiring exogenous FZD_4_ transfection (Fig. 1f). To further confirm the induction of WNT/β-catenin signaling, expression of the WNT target genes *Axin2, Lef1* and *Mfsd2a^43–45^* as well as *Mki67* were assessed. L6-F4-2 significantly enhanced these mRNAs in bEND.3 cells (Fig. 1g) as well as *Apcdd1* and *Cldn5* in cultured primary mouse brain endothelium (Supplementary Fig. 1).

### L6-F4-2 treatment rescues retinopathy in *Ndp^KO^* mice

Norrin uniquely binds the FZD_4_ receptor, which is essential for neonatal retinal vascular development, to activate WNT/β-catenin signaling^21^. In humans, mutation of *Ndp*, encoding Norrin, causes Norrie disease with congenital blindness^46^. *Ndp^KO^* mice exhibit severely impaired neonatal retinal angiogenesis with lack of capillary networks^23^, defective EC proliferation and abnormal artery-vein crossings^18^. Similarly, disruption of Norrin/FZD_4_ signaling by conditional FZD_4_ knockout in ECs reduces neonatal retinal vascularization^24^. We thus tested if our L6-F4-2 FZD_4_-selective WNT surrogate could substitute for the absence of Norrin in *Ndp^KO^* mice and rescue the mutant retina vasculature phenotype. We administered L6-F4-2 or PBS by single intravitreal (IVT) injection into the left eye at postnatal day 0 (P0) and used the right eye as a non-treatment control. P8 retinas were harvested, and veins were stained with isolectin GS-B4, revealing that PBS-injected or non-injected mice displayed characteristic *Ndp^KO^* phenotypes with reduced vein distance from optic nerve, enlarged vein diameter and loss of fine capillary networks. Notably, these impaired vein vasculatures were completely rescued by L6-F4-2 treatment of *Ndp^KO^* mice, which then resembled *Ndp* WT retinal veins (Fig. 2a, b).

**Fig. 2.**
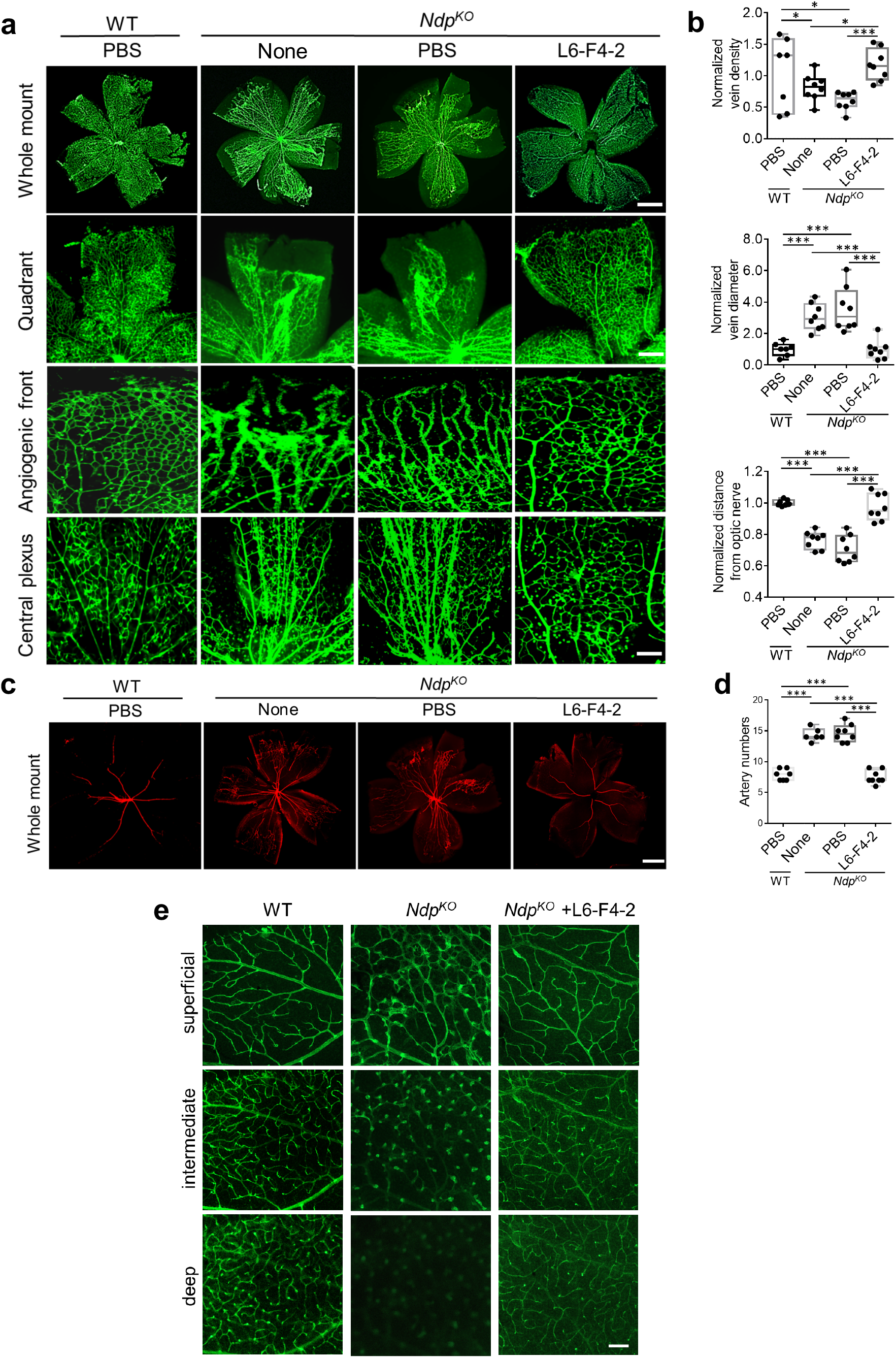
L6-F4-2 treatment rescues retinopathy in *Ndp^KO^* mice. **a-d** WT (control) and *Ndp^KO^* mice were treated with PBS or L6-F4-2 by single intravitreal injection (0.19 μg) at P0 and retinas were harvested at P8. **a** Whole-mount isolectin B4 labeling of retinal veins. Representative images of the whole retinal vascular plexus, quadrant vascular plexus, angiogenic front and central plexus are shown. Scale bars represent 0.5 mm (top), 0.3 mm (middle) and 0.15 mm (bottom two rows). **b** Quantification of vein diameter, vein density and distance from the optic nerve. **c** and **d** Wholemount anti-smooth muscle actin labeling of retinal arteries at P8 and quantification of artery branches. Scale bar represents 0.5 mm. In all the quantifications, error bars represent mean ± s.e.m., *Ndp* WT n = 7, *Ndp^KO^* n=8, *p < 0.05, ***p<0.001; Mann-Whitney U test. **e** WT (control) and *Ndp^KO^* mice were treated with PBS or L6-F4-2 by intravitreal injection (0.19 μg) at P5 and every three days with retina harvest at P21. Optical sections of isolectin-stained P21 retinas showing the three-layered retinal vasculature (superficial, intermediate, and deep layers) to document vascular architecture. Scale bars represent 100 μm.

In contrast to severely atretic retinal vein development, *Ndp^KO^* mice also manifest increased neonatal retinal arteries^18^. Whole mount arterial staining of neonatal retinas with anti-smooth muscle actin revealed that single IVT injection of L6-F4-2 restored the increased arterial phenotype of *Ndp^KO^* mice to *Ndp* WT levels (Fig. 2c, d). Optical sections were used to document malformation of the three-layered retinal vasculature. Mice injected with control antibody displayed characteristic phenotypes of *Ndp^KO^* mice including absence of intraretinal capillaries in the deep layer and glomeruloid vascular malformations instead of a properly formed capillaries in the intermediate layer. Angiogenesis was virtually restored and intermediate and deep intraretinal capillary beds were properly formed in *Ndp^KO^* mice injected with L6-F4-2 (Fig. 2e).

In WT retinas, major arteries and veins are segregated into alternating and radially arrayed territories, and rarely cross. By contrast, P8 *Ndp^KO^* retinas showed an average of 4 crossings of major arteries and veins and many more crossings in smaller vessels, consistent with prior reports^18^, which was then strongly reduced by L6-F4-2 versus PBS or no injection (Supplementary Fig. 2a, b). *Ndp^KO^* retinal vessels have been reported to exhibit increased filopodia^47^. At P8, *Ndp^KO^* retinas displayed elevated filopodia, and decreased Ki67+ EC proliferation, vein density and branch points, all of which were rescued by L6-F4-2 treatment (Supplementary Fig. 2c-h). Notably, single IVT injection of tetravalent WNT surrogates recognizing FZD_1/2/7_ or FZD_5/8_ at P0 did not rescue *Ndp^KO^* retinal vein or artery phenotypes at P8, confirming the striking specific efficacy of the FZD_4_-selective L6-F4-2 agent (Supplementary Fig. 3).

### L6-F4-2 activates retinal endothelial WNT/β-catenin signaling

We next examined L6-F4-2 action by bulk RNA-seq of retinal endothelium from PBS or L6-F4-2-treated WT versus *Ndp^KO^* mice, after single IVT injection at P0, retina harvest at P8 and FACS isolation of CD31+ ECs. This bulk RNA-seq identified retinal EC mRNAs corresponding to WNT target genes that were decreased by *Ndp^KO^* mice but re-induced by L6-F4-2 (Fig. 3a, b), consistent with the diminution of WNT/β-catenin signaling activity in *Ndp^KO^* mice^10^ and the rescue of *Ndp^KO^* developmental retinal angiogenesis by L6-F4-2 (Fig. 2). This *Ndp^KO^*- and L6-F4-2-regulated gene set overlapped with blood vessel development genes (GO:001568) and numerous established WNT/β-catenin signaling mRNAs (*Apcdd1*, *Axin2*, *Tcf7*, *Celsr1*; GO:0016055) including those known to be WNT-regulated in BBB ECs^15,48^ (Fig. 3b). We re-validated these findings by qRT-PCR of CD31 + retinal ECs, indicating that the WNT downstream target genes, *Axin2* and *Apcdd1* and blood vessel development-related genes *Tbx1* and *Tgfa* were all reduced in *Ndp^KO^* mice but rescued by L6-F4-2 (Fig. 3c).

**Fig. 3.**
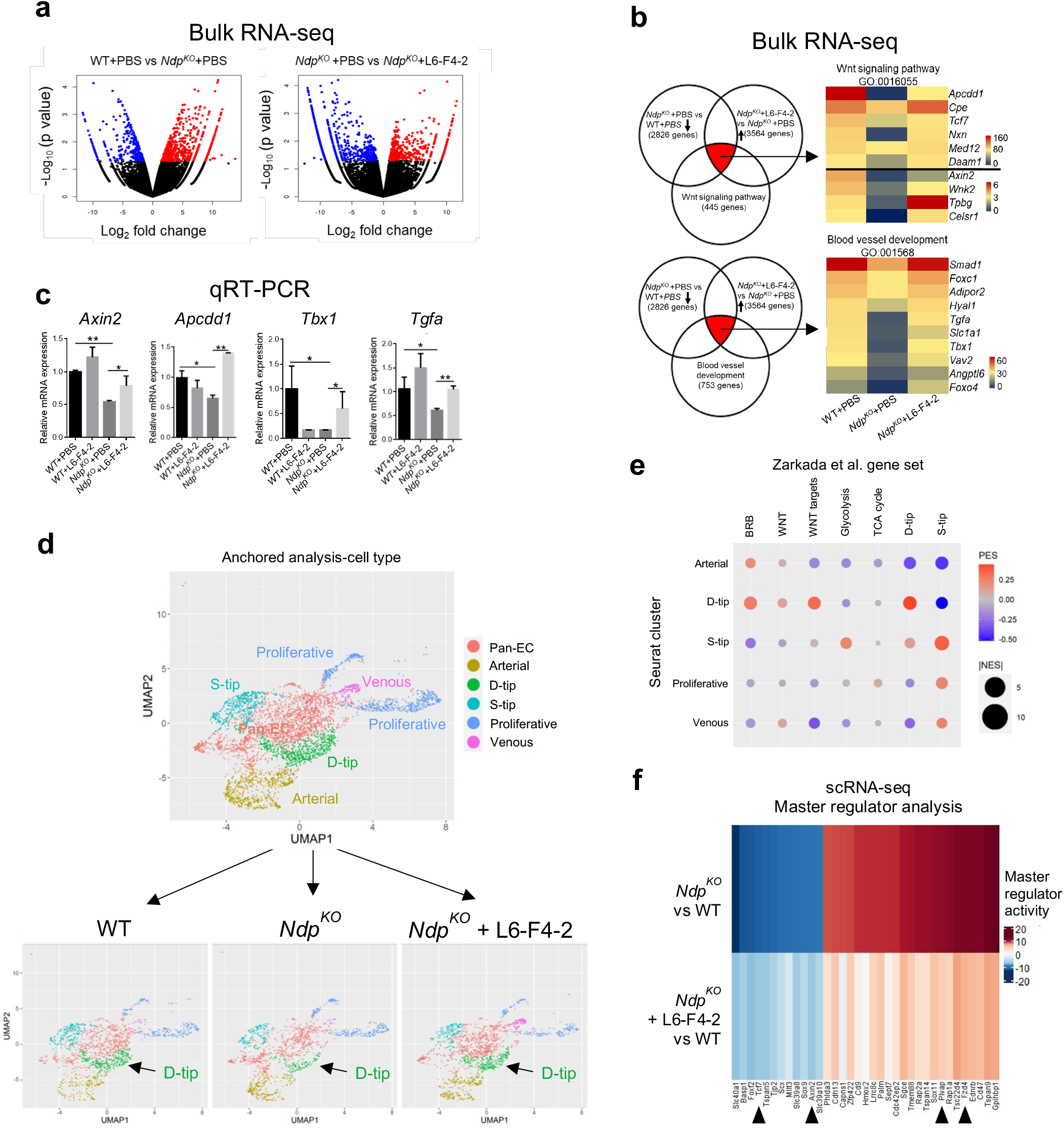
Effects of L6-F4-2 on retinal endothelial WNT/β-catenin signaling and vascular subtypes. **a-c** WT (control) and *Ndp^KO^* mice were treated with PBS or L6-F4-2 by intravitreal injection (0.19 μg) at P0 and 5-7 pooled retinas for each group were harvested at P8. Retinal vessel ECs were purified by anti-CD31 FACS and ultra-low input bulk RNA-seq analysis was performed. **a** Left: WT (PBS-treated) vs *Ndp^KO^* (PBS-treated). Right: *Ndp^KO^* (PBS-treated) vs *Ndp^KO^* (treated with L6-F4-2). Volcano plot depicting genes with a p-value < 0.05 and log_2_ fold-change > 1 are indicated by red dots which represent upregulated genes. Genes with a p-value < 0.05 and log_2_ fold change < −1 are indicated by blue dots, representing downregulated genes. **b** Left: Venn diagrams showing overlap in bulk RNA-seq differentially expressed genes following treatment with PBS/L6-F4-2 in WT or *Ndp^K**O**^* mice. Right: Heat maps of the top 10 expressed genes mapping to the indicated WNT/β-catenin signaling pathway and blood vessel development biological processes across treatment groups in the RNA-seq experiment. **c** qRT-PCR validation of differentially expressed genes identified by RNA-seq. Error bars represent mean ± s.e.m., n=3, *p < 0.05, **p<0.01; Mann-Whitney U test. **d-f** Single cell RNA-seq with the same treatment conditions as (**a-c**). **d** scRNA-seq UMAP plots of retinal EC sub-clusters from P8 retinas from all conditions merged (top) and from the WT, *Ndp^KO^*, and *Ndp^KO^* + L6-F4-2 conditions (bottom). Alteration of the D-tip population is indicated by arrows. **e** Dot plot of NaRnEA enrichment for previously described Zarkada et al. gene sets^49^. Dots indicate the enrichment of specified gene sets from ref ^49^ (x-axis) in a given Seurat cluster (y-axis). Dot size is determined by the absolute value of the normalized enrichment score (NES); large dots indicate more significant enrichment. Dot color is determined by the proportional enrichment score (PES), which indicates the sign of the enrichment. Enrichment analysis was performed with the NaRnEA algorithm. BRB, brain-retinal-barrier. **f** Master regulators of the *Ndp^KO^* condition versus WT as identified by PISCES analysis of the scRNA-seq data with reduced or reverted activity profiles in the *Ndp^KO^* + L6-F4-2 condition. These proteins represent transcriptional programs that have been reverted by the therapeutic agent.

### L6-F4-2 rescue of endothelial D-tip cells in *Ndp^KO^* retina

We then performed single cell RNA sequencing (scRNA-seq) on FACS-purified CD31+ P8 retinal ECs from *Ndp^KO^* versus WT mice after P0 IVT injection of L6-F4-2 or PBS. This scRNA-seq data was analyzed for changes in endothelial subsets between the WT, *Ndp^KO^*, and *Ndp^KO^* + L6-F4-2 treatment conditions. We first used the Seurat pipeline to anchor the retinal endothelial scRNA-seq data from the three experimental conditions, focusing on the >95% of sorted cells expressing the pan-endothelial markers *Cdh5 and Pecam1* to generate cell clusters in UMAP space. Prior single cell RNA-seq has defined several neonatal retinal vascular cell types, including two classes of tip cells, which remain superficially (S-tip) on the retinal surface or dive (D-tip) into the neural retina respectively, as well as capillary, venous, arterial, and proliferative populations^49^. We thus applied the markers from this prior retinal endothelial classification^49^ to assign cell type identities to our unsupervised scRNA-seq Seurat analysis (Fig. 3d and Supplementary Fig. 4). This identified *Cldn5^+^Mfsd2*^+^*Spock2*^+^ D-tip, *Angpt2*^+^*Esm1*^+^ S-tip, *Unc5b*^+^*Bmx*^+^ arterial, *Ptgis^hgh^* venous and two clusters of *Mki67*^+^*Birc5*^+^ proliferative endothelium (Fig. 3d, Supplementary Fig. 4). The remaining cells expressed *Cdh5 and Pecam1* but were not enriched for markers of other clusters or for previously described capillary markers^49^ and therefore were described as pan-endothelial (Pan-EC) (Fig. 3d, Supplementary Fig. 4). Additional gene set variation analysis (GSVA) using NaRnEA^50^ demonstrated strong correlation between GSEA processes in from D-tip and S-tip clusters from prior datasets^49^ compared to our scRNA-seq analysis, further confirming these cell type assignments. Besides this global overlap, the strongest associations for Wnt signaling, glycolytic processes and TCA cycle were with D-tip, S-tip and proliferative subsets respectively (Fig. 3e), consistent with prior reports^49^. Our studies did not allow high-confidence identification of previously described capillary populations^49^, which may reside in the pan-EC cluster (Fig. 3d, Supplementary Fig. 4).

We focused on changes in cell type frequency between the identified populations in WT, *Ndp^KO^* and *Ndp^KO^* + L6-F4-2 using a chi-squared test, leveraging standardized residuals to identify significantly altered populations. By far, the strongest and most dominant effect between WT and *Ndp^KO^* was significant depletion in D-tip cells (p = 7.4 × 10^-13^), which was subsequently rescued by L6-F4-2 (p = 1.2 × 10^-8^) (Fig. 3d). These changes in D-tip endothelium were notably consistent with (i) the profoundly decreased vascularization of deep retinal layers in *Ndp^KO^* and subsequent rescue by L6-F4-2 in vivo (Fig. 2d) and (ii) the finding that D-tip cells were also the subset having the strongest enrichment in WNT target genes (Fig. 3e). S-tip cells were less significantly decreased in *Ndp^KO^* (p = 0.002) and not rescued by L6-F4-2 (p = 0.44) (Fig. 3d), potentially consistent with similar numbers of superficial endothelium, albeit with differential radial extension, in these three conditions (Fig. 2e).

### Master regulator analysis of L6-F4-2 action in retinal vasculature

We further stratified the scRNA-seq data into transcriptional nodes using master regulator analysis. A gene expression signature was generated for *Ndp^KO^* v. WT and *Ndp^KO^* + L6-F4-2 v. WT using the DESeq2 package in R. We then applied the PISCES (Protein Activity Inference in Single Cells) pipeline, to first generate a transcriptional network using ARACNe-AP^51^ (Algorithm for the Reconstruction of Accurate Cellular Networks with Adaptive Partitioning) from the WT data with k = 5 metacells, and then inferred protein activity using VIPER^52^ (Virtual Inference of Protein Activity by Enriched Regulon Analysis) to detect concomitant changes in downstream target mRNAs. The top such inferred proteins in each scRNA-seq experimental condition were selected as candidate master regulators (MRs), and these were compared to identify which significant retinal endothelial MRs in the *Ndp^KO^* condition reverted to a more WT-like state upon L6-F4-2 rescue (*Ndp^KO^* + L6-F4-2). QC metrics indicated robust library preparation (Supplementary Fig. 5), and the data were filtered based on thresholds in Supplementary Table 1.

The most downregulated MR activities in *Ndp^KO^* mice (left column, deep blue) were substantially rescued by L6-F4-2 treatment of *Ndp^KO^* mice (left column, light blue) (Fig. 3f). Indeed, this unbiased analysis again revealed *Axin2* and *Tcf7* as MRs decreased in *Ndp^KO^* but re-induced by L6-F4-2 (Fig. 3d), identical to bulk RNA-seq and qRT-PCR data (Fig. 3a-c). Plasmalemma vesicle-associated protein (PLVAP) is an EC fenestration component that regulates BBB/BRB permeability and is upregulated in pathological conditions associated with compromised barrier function and is decreased by WNT/β-catenin signaling^48,53,54^. Accordingly, *Plvap* was an inferred MR that was indeed overexpressed in *Ndp^KO^* versus WT mice (Fig. 3f, right column, red), but downregulated by superimposed L6-F4-2 treatment (Fig. 3f, right column, pink). Interestingly, the MR *Fzd4* was increased in *Ndp^KO^* but suppressed by L6-F4-2, potentially consistent with a feedback loop where absence of the Norrin ligand elicits compensatory upregulation of the receptor FZD_4_ and its signaling network, which is rescued by L6-F4-2 (Fig. 3f, right column). We also identified MRs which were substantially dysregulated instead of rescued by L6-F4-2 (Supplementary Fig. 6).

### L6-F4-2 treatment promotes BBB and BRB function in neonatal and young mice

Norrin/FZD_4_/β-catenin signaling is required for the development and maintenance of CNS endothelial cell barrier properties. Constitutive KO of *Ndp or* FZD_4_ elicits BBB permeability in cerebellum, which can be rescued by EC-specific transgenic expression of stabilized β-catenin or transgenic restoration of Norrin expression^11,18^. Further, FZD_4_ conditional KO in retinal ECs induces BRB leak^11,18^. We thus asked if L6-F4-2 could represent a novel therapeutic agent for CNS endothelial barrier dysfunction. Pharmacokinetic analysis of L6-F4-2 after i.p. or i.v. administration for bioavailability and clearance information suggested an extended half-life for L6-F4-2 of approximately 3 days, as expected of an antibody molecule (Supplementary Fig. 7).

We then performed i.p. injection of L6-F4-6 or PBS into *Ndp* WT and KO mice at P0, P7 and P14 with harvest at P21. As expected, Sulfo-NHS-biotin did not extravasate from the retinal vasculature of postnatal WT mice, consistent with an intact BRB. In contrast, *Ndp^KO^* retina exhibited substantial extravascular biotin leak, in agreement with published phenotypes^11,18^. However, L6-F4-2 treatment strongly improved BRB function, reducing biotin extravasation in *Ndp^KO^* retina to essentially WT levels (Fig. 4a, b). *Ndp^KO^* cerebral cortex did not exhibit biotin extravasation because of an intact BBB. Extensive vascular leak was observed in the parenchyma of WT and *Ndp^KO^* liver and kidney, organs with fenestrated vasculature, which as expected was not significantly reversed by L6-F4-2 (Fig. 4a, b).

**Fig. 4.**
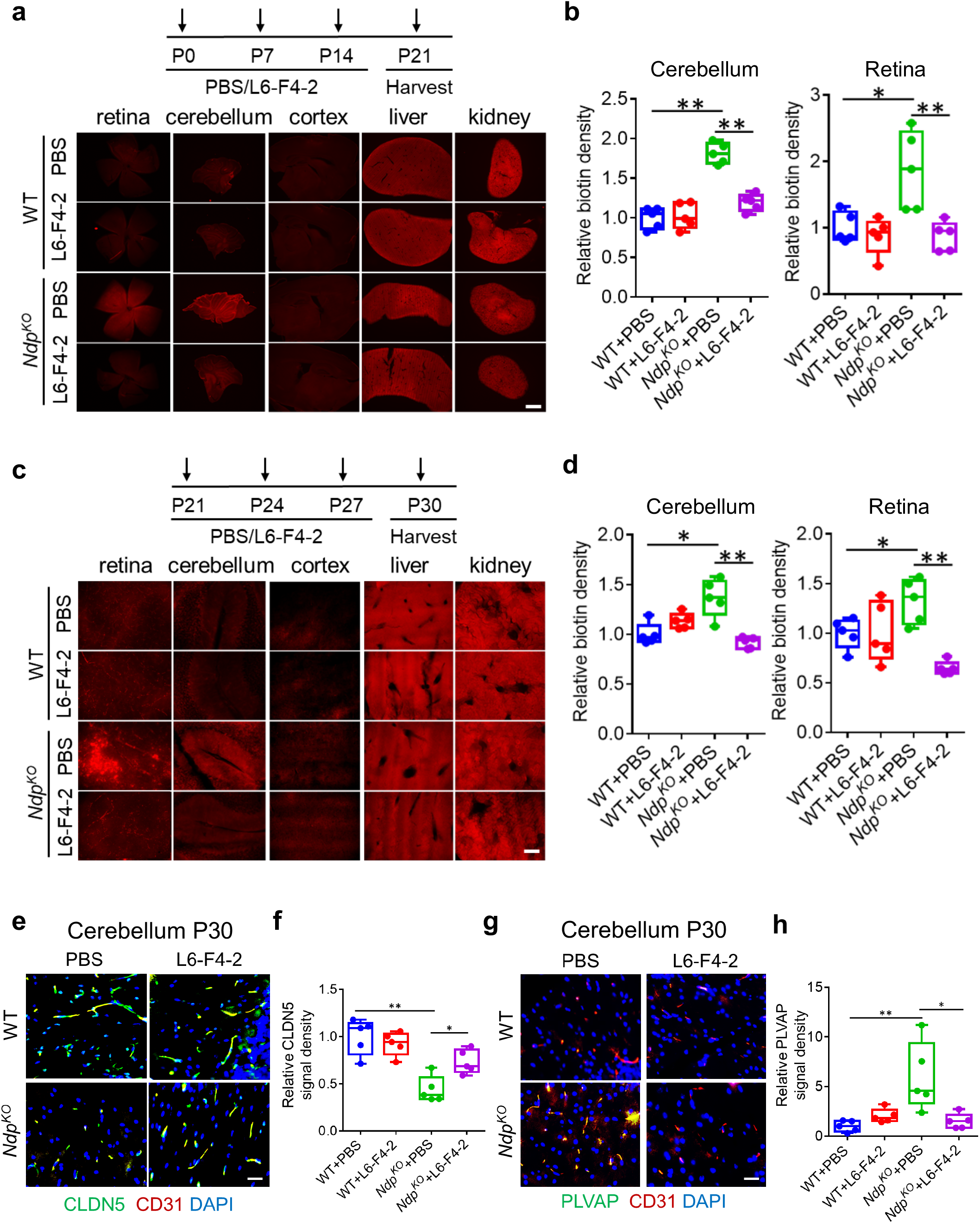
L6-F4-2 treatment promotes endothelial blood-brain and blood-retina barrier function. **a** WT (control) and *Ndp^KO^* mice were treated with PBS or L6-F4-2 by i.p. injection (2.5 mg/kg) at P0, P7 and P14 and tissues were harvested at P21. Sulfo-NHS-biotin staining reveals BBB/BRB defects in cerebellum and retina, but not cerebral cortex in *Ndp^KO^* mice, while these defects can be rescued by L6-F4-2 treatment. Biotin labeling of liver and kidney parenchyma serve as a positive control. Scale bar, 500 μm. **b** Quantification of (**a**) for BBB/BRB leakage in *Ndp^KO^* cerebellum and retina with rescue by L6-F4-2 treatment. **c** Mice were treated at P21, P24 and P27 (2.5 mg/kg, i.p.) and tissues were harvested at P30. Sulfo-NHS-biotin staining reveals BBB/BRB defects in cerebellum and retina, but not cerebral cortex in *Ndp^KO^* mice with rescue by L6-F4-2. Scale bar, 200 μm. **d** Quantification of (**c**) with BBB/BRB leakage in *Ndp^KO^* cerebellum and retina and reversal by L6-F4-2. **e** L6-F4-2 rescues barrier function defects in P30 *Ndp^KO^* mice with increased expression of the tight junction component CLDN5, P30 cerebellum IF, overlay of CD31 IF and DAPI. **f** Quantification of (**e**). **g** L6-F4-2 decreases expression of the EC fenestration component PLVAP in P30 *Ndp^KO^* mice, with overlay of CD31 IF and DAPI. **h** Quantitation of (**g**). For e-h, scale bars represent 100 μm. To quantify CLDN5 and PLVAP in (**f**, **h**), the density was measured with ImageJ and normalized to vessel area (CD31). Error bars represent mean ± s.e.m., n=5, *p < 0.05, **p < 0.01; Mann-Whitney U test.

We studied whether L6-F4-2 could restore *Ndp^KO^* BBB and BRB defects in more aged mice by administering L6-F4-2 at P21, P24, P27 and harvesting tissues at P30. L6-F4-2 enhanced BRB and BBB barrier integrity in *Ndp^KO^* mice, as quantification of biotin density in both cerebellum and retina revealed significant rescue by L6-F4-2 versus PBS (Fig. 4c, d). In WT mice, the cerebral cortex exhibited an intact BBB while the fenestrated liver and kidney endothelium again manifested constitutive vascular leakage that was L6-F4-2-insensitive (Fig. 4c). The strong BBB defects in *Ndp^KO^* mice were appropriately associated with reduced endothelial CLDN5 and increased PLVAP expression. However, L6-F4-2 treatment of *Ndp^KO^* mice strongly upregulated CLDN5 (Fig 4e, f) and repressed PLVAP in endothelium of brain (Fig. 4g, h) and retina (Supplementary Fig. 8), consistent with restoration of WNT/β-catenin signaling and barrier maturation. In total, these results indicated that L6-F4-2 not only reversed BRB phenotypes but also was robustly active in the BBB, indicating broad utility of FZD_4_ agonism.

### L6-F4-2 treatment rescues stroke phenotypes in adult WT mice

To expand the potential applications of FZD_4_-selective WNT surrogates beyond *Ndp^KO^* mice to prevalent medical conditions, we investigated the effect of L6-F4-2 in adult ischemic stroke. Despite the substantial incidence of stroke-related death and disability^55^, treatment options are limited to mechanical stenting or pharmacologic therapy with tissue plasminogen activator (tPA), albeit with limited temporal windows and hemorrhage risk^56–58^. Wild-type adult C57Bl/6J mice (6-8 weeks) were subjected to transient middle cerebral artery occlusion stroke (tMCAO) by catheter occlusion to induce cerebral ischemia for 45 minutes, followed by reperfusion for 2 days. L6-F4-2 (3 mg/kg, i.v.) or control NIST mAb (National Institute of Standards and Technology, humanized IgG1κ monoclonal antibody) were administered at 1 and 24 hours after tMCAO. Brains were harvested 48 hours post-stroke and infarct TTC staining was immediately performed after tissue collection (Fig. 5a). The NIST and L6-F4-2 arms contained 20 mice each, of which n=13 and n=18 were evaluable after 48 h because of poststroke mortality. Notably, L6-F4-2 substantially decreased cerebral infarct volume versus control mice (control n=13, L6-F4-2 n=18, p<0.01) (Fig. 5b, c). Stroke-induced BBB permeability was assessed by IgG staining as a measure of plasma protein leak into the brain parenchyma. Accordingly, L6-F4-2 strongly reduced cerebral IgG extravasation versus NIST controls (control n=13, L6-F4-2 n=18, p<0.05), consistent with BBB functional rescue (Fig. 5b, d). In addition, we assessed BBB integrity by Sulfo-NHS-biotin tracer extravasation. Strikingly, L6-F4-2 treatment significantly alleviated the leakage of biotin in stroke regions and no significant differences were observed in non-stroke areas (Fig. 5e, f). The L6-F4-2-induced rescue of infarct size and BBB integrity was further paralleled by significant reduction in edema (p<0.05) (Fig. 5g) and neurological score (p=0.0312) (Fig. 5h), compared against NIST treatment.

**Fig. 5.**
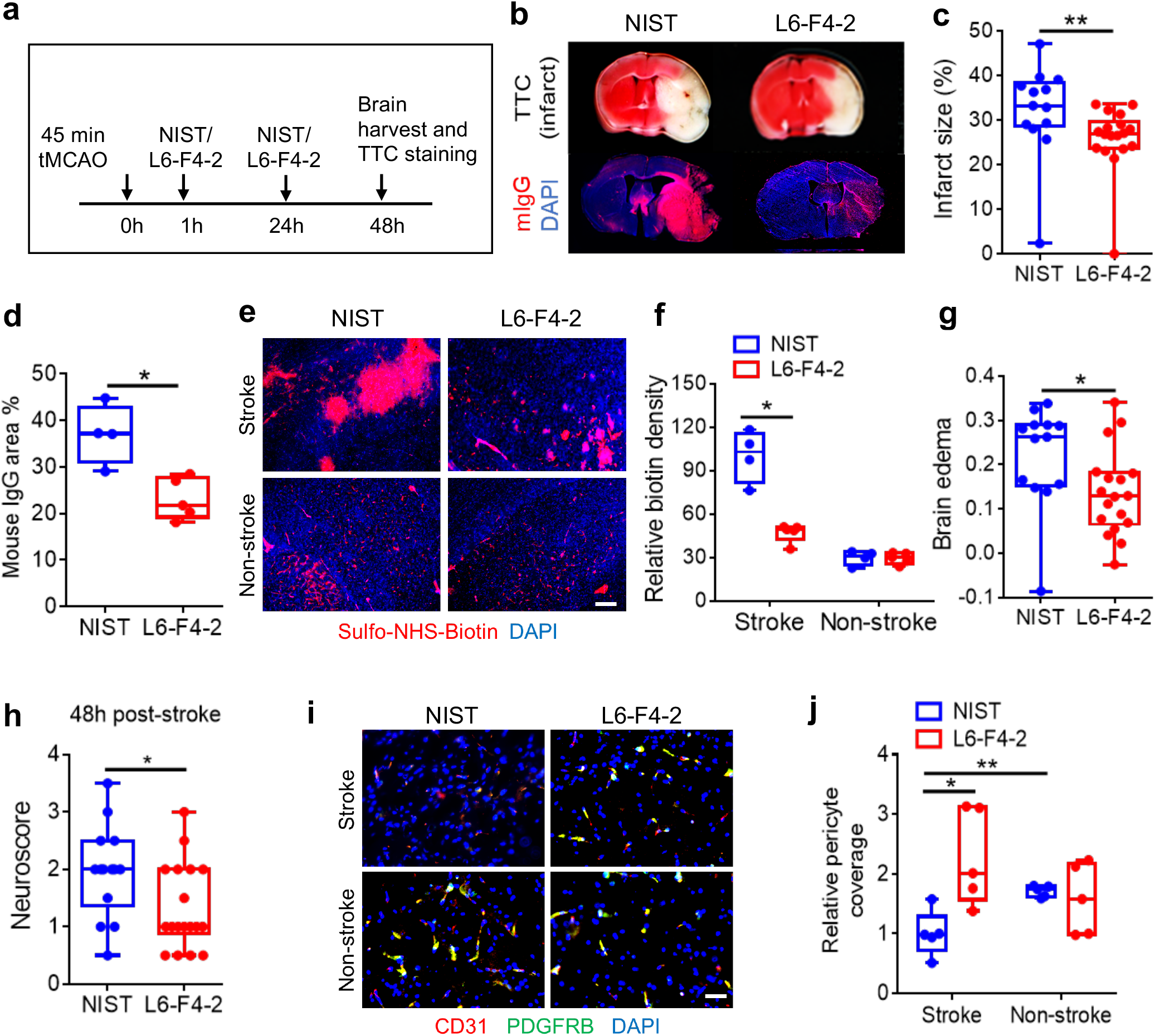
L6-F4-2 treatment rescues stroke phenotypes in wild-type mice. **a** Schematic of tMCAO surgery with 45 min occlusion time, L6-F4-2 treatment time course and brain harvest at day 2 post-stroke. **b** TTC staining (top), and mouse IgG extravasation (mIgG) (bottom) of coronal sections at 48 hours post-stoke. **c** Quantification of infarct size for NIST control mAb (n=13) and L6-F4-2 (n=18). **d** Quantification of mouse IgG staining, NIST control n=4, L6-F4-2 n=5. **e** Representative images of BBB integrity in brains of the indicated mice in stroke and non-stroke regions after 45 min tMCAO, 2 days of reperfusion and two L6-F4-2 treatments (3 mg/kg, i.v.), as assessed by the Sulfo-NHS-biotin tracer extravasation assay. Scale bar, 50 μm. **f** Quantification of extravasated exogenous tracer Sulfo-NHS-biotin using ImageJ. n = 5. **g** Fractional change in brain edema for NIST control mAb (n=13) and L6-F4-2 (n=18). **h** Neurological scores at 48 h after tMCAO surgery for NIST control mAb (n=13) and L6-F4-2 (n=18). **i** Co-immunofluorescence staining for PDGFRB and CD31 in infarcted brain (stroke and non-stroke) regions and **j** quantification of pericyte coverage. The PDGFRB signal was normalized to CD31; n = 5. Scale bar, 100 μm. Error bars represent mean ± s.e.m., *p < 0.05, **p <0.01, Mann-Whitney U test.

Post-stroke effects of L6-F4-2 on neurovascular unit endothelial cells and pericytes, both of which are essential for BBB integrity, were characterized by IF staining of stroke and non-stroke hemispheres. Pericyte coverage of brain endothelium is markedly reduced post-stroke^15,59-61^ and the decreased pericyte coverage in the stroke hemisphere was strongly restored by L6-F4-2 treatment. In contrast, the contralateral non-stroke hemisphere in control and L6-F4-2 treated groups did not significantly differ in their substantial pericyte coverage of the endothelium (Fig. 5i, j).

## Discussion

The specialized blood-brain and blood-retinal barriers stringently regulate entry of substances from the circulation into the CNS parenchyma. The essential nature of these barrier functions is exposed by edema, hemorrhage and inflammation during diverse CNS pathologies including neoplasia, neural degeneration, autoimmunity and infection^3–6^. FZD_4_ is an attractive therapeutic target given its high expression in CNS endothelium^15,19,62^, essential functions in the BRB and cerebellar BBB, and functional redundancy with GPR124/RECK/WNT7 in cortical BBB regulation^10,11,18^. However, therapeutic use of FZD_4_ ligands is complicated by palmitoylation and insolubility of WNTs and difficulties in expressing Norrin^33,34^. GPR124-selective WNT7 mutants have been described but are predicted to retain palmitoylation, and are thus poorly compatible with systemic administration, therefore requiring local viral vector delivery within the BBB^63^.

Here, we describe development of L6-F4-2, a selective tetravalent potent surrogate of the FZD_4_:LRP6 receptor complex. L6-F4-2 does not bind FZD_1-3_ or FZD_5-10_ ECDs but promotes proximity of FZD_4_ to LRP6 coreceptors, consistent with our prior synthesis of bioengineered WNT surrogates targeting other FZD family members^35–38,40,64^. Compared to Norrin and the recently reported FZD_4_-selective WNT surrogate F4L5.13^28^, L6-F4-2 exhibits approximately 100 times higher FZD_4_ affinity and potency of WNT/β-catenin signaling activation in vitro. As opposed to the obligate palmitoylation of highly hydrophobic native WNTs, L6-F4-2 is non-lipidated, facilitating systemic delivery, efficacy, as well as ease of production. At the same time, FZD_4_ tropism may have further advantages of averting promiscuous activation of other FZD family members.

In vivo efficacy of L6-F4-2 is strongly supported by robust rescue of retinal angiogenesis and BRB leakage in *Ndp^KO^* newborn pups, where L6-F4-2 substitutes for the absent Norrin. In the *Ndp^KO^* model, L6-F4-2 induces WNT-dependent gene expression in retinal endothelium, consistent with an on-target mechanism. Our scRNA-seq analysis revealed D-tip cells as the most significantly dysregulated endothelial cell type in *Ndp^KO^* retina, which was indeed potently rescued by L6-F4-2 treatment. These findings are consonant with prior studies indicating particular enrichment of WNT signaling processes within D-tip cells^49^. While L6-F4-2 restored BRB function at both P21 and P30, neither angiogenesis deficits nor functional visual acuity were substantially reversed at P30, indicating that at late time points fixed capillary patterning defects supervene that are L6-F4-2-refractory, contrasting with the reversibility of barrier phenotypes (JD and CJK, unpublished data). We utilized single intravitreal injection of L6-F4-2, analogous to intravitreal administration of VEGF inhibitors for macular degeneration and diabetic retinopathy^65^, in contrast to systemic treatment of retinal phenotypes with FZD_4_-selective WNT surrogates such as F4L5.13^28^.

Lastly, our present studies are notable for substantially extending the therapeutic utility of FZD_4_ agonism. Prior efforts have been restricted to studying lower affinity FZD_4_-selective WNT surrogates in the retina of knockout mice lacking *Tspan12*, encoding a Norrin co-receptor^28^, or rescue of *Ndp^KO^* retinopathy by transgenic expression of Norrin^18^. Crucially, our results with L6-F4-2 extend the successful demonstration of FZD_4_ agonism from the BRB to additional indications in the blood-brain barrier, as evidenced by robust rescue of *Ndp^KO^* cerebellar BBB permeability. Notably, L6-F4-2 also markedly improved stroke outcomes and BBB function in wild-type adult C57Bl/6J mice, thus enlarging potential benefits beyond mutations in *Ndp/Tspan12*, to prevalent and disabling medical conditions in the general population such as cerebral infarction. Indeed, FZD_4_ agonism may certainly benefit Mendelian disorders such as Norrie disease or other WNT pathway-mutant hereditary retinopathies, as facilitated by our demonstration of successful intravitreal delivery. However, our results suggest that FZD_4_-selective WNT surrogates could find broader general application to conditions ranging from stroke, neurodegeneration, neoplasia, autoimmunity, and infection, representing neurological diseases at-large characterized by disrupted BBB function.

## Acknowledgements

We are grateful to members of the Kuo laboratory for constructive discussions. We acknowledge Rachel Lam, Nay Saw and Mehrdad Shamloo for conducting tMCAO experiments in the Stanford University Behavioral and Functional Neuroscience Laboratory. We thank Ming Chen and Hannah Schmitz from the Department of Genetics, Stanford University School of Medicine, for technical assistance with IVT injections, and Jay Tibbitts from Surrozen for assistance with PK analysis.

## Author contributions

J.D., S.L., W.C.Y., Y.L. and C.J.K. conceived the project. S.L. designed and performed in vitro experiments related to L6-F4-2. J.D. designed and conducted in vitro, and animal experiments related to retina and stroke studies. L.V. and V.V.U. analyzed bulk and single cell RNA sequencing data. K.Y. performed FACS analysis and helped with animal experiments. C.C.R. conducted and guided statistical analysis of retina vascular architecture rescue experiments. M.V. contributed to assay results in Figure 1. H.C. and A.S. contributed to the construction, expression, and purification of L6-F4-2. H.N and W.N. conducted PK studies. A.K.M., M.T. and S.A. conducted sample processing and technical support. L. Kim and L. Kelly performed mouse IVT injections. D.V., A.J.C., Y.L., W.C.Y. and C.J.K. participated in data analysis and interpretation. J.D., S.L., L.V., C.C.R., Y.L. and C.J.K. wrote the manuscript. All authors reviewed and edited and approved the final manuscript.

## Data and materials availability

All data are available in the main text or the supplementary materials and upon request.

## Methods

### Molecular cloning and protein generation

Constructs for F4-2_Fab (hereafter F4-2) and L6-F4-2 were cloned into the pcDNA3.1(+) mammalian expression vector (Thermo Fisher). The F4-2 heavy chain contains a 6xHis tag on its C-terminus. For the tetravalent constructs for L6-F4-2, each light chain variable region (VL) or heavy chain variable region (VH) of YW211.31.57 was fused onto the either light chain or heavy chain N-terminus of the F4-2 IgG through a 15-mer linker (GGGGSGGGGSGGGGS), which contains L234A/L235A/P329G mutations (LALAPG) to eliminate effector function^66^.

Recombinant proteins were produced in Expi293F cells (Thermo Fisher Scientific) by transient transfection. The F4-2_Fab protein was first purified using cOmplete® His-tag purification resin (Sigma-Aldrich), and L6-F4-2 protein was first purified with CaptivA Protein A affinity resin (Repligen). All proteins were further polished with Superdex 200 Increase 10/300 GL (GE Healthcare Life Sciences) size-exclusion chromatography (SEC) using 1 × HBS buffer (20 mM HEPES pH 7.4, 150 mM NaCl) or 2xHBS buffer (40 mM HEPES pH 7.4, 300 mM NaCl). The CRDs of 10 FZDs were expressed and purified as previously described^40^. All proteins were also examined by SDS-polyacrylamide electrophoresis and estimated to be > 90% purity.

### Affinity measurement and binding specificity

Binding kinetics of F4-2_Fab or L6-F4-2 to FZD CRDs were determined by bio-layer interferometry (BLI) using an Octet Red 96 (PALL ForteBio, Fremont, CA) instrument at 30°C, 1000 rpm with streptavidin (SA) biosensors. Biotinylated FZD_4_ CRD was diluted to 50 nM in the running buffer (PBS, 0.05% Tween-20, 0.5% BSA, pH 7.2) and captured to the SA biosensor followed by dipping into wells containing the F4-2_Fab or L6-F4-2 at different concentrations in running buffer or into a well with only running buffer as a reference channel. The dissociation of the interaction was followed with the running buffer. K^D^ for each binder was calculated by Octet System software, based on fitting to a 1:1 binding model. For binding specificity test, L6-F4-2 was diluted to 50 nM in the running buffer and captured to the AHC biosensor followed by dipping into wells containing 200 nM FZD CRDs in running buffer. The dissociation of the interaction was followed with the running buffer.

### SuperTop Flash (STF) assay

WNT/β-catenin signaling activity was measured using HEK293, HRMEC, or bEnd.3 cells containing a luciferase gene controlled by a WNT-responsive promoter (Super Top Flash reporter assay, STF) as previously reported^36^. In brief, cells were seeded at a density of 10,000 per well in 96-well plates 24 hours prior to treatment at the presence of 3 μM IWP2 to inhibit the production of endogenous WNTs. L6-F4-2 proteins were then added to the cells overnight. Recombinant human WNT3A (R&D Systems) and recombinant human Norrin (R&D Systems) were used as response comparators. Cells were lysed with Luciferase Cell Culture Lysis Reagent (Promega) and luciferase activity was measured with Luciferase Assay System (Promega) using vendor procedures.

### Purification and culture of mouse brain endothelial cells

Mouse brain endothelial cells were isolated from the cortex of adult C57BL/6J mice by established procedures followed by in vitro puromycin selection and in vitro culture in EGM-2MV as described^15^.

### Quantitative PCR analysis of gene expression

Either bEnd.3 cells or primary mouse brain endothelial cells were treated with 10 nM L6-F4-2 or NIST mAb for 24 h, followed by RNA extraction using the Qiagen RNeasy Micro Kit (Qiagen, Hilden, Germany) or the MagMAX mirVana Total RNA Isolation Kit (ThermoFisher, A27828), respectively. cDNA was produced using the SuperScript IV VILO cDNA Synthesis Kit (ThermoFisher, Waltham, MA). RNA was quantified using TaqMan Fast Advanced Master Mix with the following probes Mm00443610_m1 *Axin2*, Mm01278617_m1 *Mki67*, Mm01192208_m1 *Mfsd2a*, Mm00550265_m1 *Lef1* (ThermoFisher, 4331182). Values were normalized to expression of constitutive *ActinB* RNA using Mm02619580_g1 (ThermoFisher, Waltham, MA).

RNA from FACS-sorted retinal endothelial cells was extracted using Arcturus PicoPure RNA Isolation Kit (Applied Biosystems) and reverse transcribed by the SuperScript IV VILO cDNA Synthesis Kit (ThermoFisher, Waltham, MA). cDNA was pre-amplified using SsoAdvanced™ PreAmp Supermix (BioRad, Hercules, CA). qPCR was performed using SYBR green Master Mix with primers (5’ à 3’) as follows: *Axin2* (Forward: GCCGACCTCAAGTGCAAACTC, Reverse: GGCTGGTGCAAAGACATAGCC); *Apcdd1* (Forward: AACCCCACCTACACCCTCATC, Reverse: CGCCGTGAAGCTGGTAGTC); *Tbx1* (Forward: CTGTGGGACGAGTTCAATCAG, Reverse: TTGTCATCTACGGGCACAAAG) and *Tgfa* (Forward: CACTCTGGGTACGTGGGTG, Reverse: CACAGGTGATAATGAGGACAGC). Relative RNA expression was normalized to *Gapdh* (Forward: TGAACGGGAAGCTCACTGG, Reverse: TCCACCACCCTGTTGCTGTA). Cldn5 primers are previously described^15^.

### Mice

*Ndp^KO^* mice (#012287) and WT mice (C57BL/6J #000664) were purchased from the Jackson Laboratory. Mice were housed and bred in a normal experimental room and exposed to a 12-hour light/dark cycle with free access to food and water. All procedures were performed in accordance with approved IACUC protocols at Stanford University.

### Intravitreal injection

For IVT injection in P0 pups, animals were anesthetized via isoflurane induction. Once anesthetized and devoid of a pedal response, a small incision ~1.5 mm was made with the tip of a surgical scalpel blade or straight vannas scissors along the eyelid margin. Curved forceps were used to poke the eye. The tip of a beveled needle was used to make a hole through the sclera into the vitreous space. A borosilicate glass tip was slid through the hole, and 0.5 μl L6-F4-2 (0.19 μg) or PBS was injected. The eye was gently pushed back into its socket, and a drop of lidocaine (2%) was administered.

### Whole mount immunofluorescent staining and imaging of mouse retina

Mice were humanely sacrificed, and eyes were harvested and transferred into 4% paraformaldehyde (PFA) to fix for 10-15 min and then to cold PBS on ice in a Petri dish for dissection. Fat surrounding the eye was displaced. The edge of the cornea was pierced with sharp scissors and cut around the cornea and iris and discard. The lens and vitreous humour were removed using forceps. The hyaloid vessels were removed by gathering them in the center of the eye with fine forceps then quickly pulled away from the center of the retina. Then, 4 to 5 radial incisions reaching approximately 2/3 of the radius of the retina were created using spring scissors to create a ‘petal’ shape. PBS was drawn off using a Pasteur pipette to flatten the retina, and then excess PBS was removed with a small piece of absorbent paper. Cold (−20 °C) methanol was slowly dropped onto the retinal surface until covered and flooded with additional methanol, transferred into 48-well plates and gently rinsed in PBS. Next, PBS was removed, and retinas were covered with 100 μl of Perm/Block solution (PBS + 0.3% Triton + 0.2% BSA) for 1 h, incubated overnight at 4°C with primary antibodies (Isolectin GS-IB4, Alexa Fluor™ 488 Conjugate or Anti-Actin, α-Smooth Muscle–Cy3™ antibody) and washed for 4 × 10 min in PBS + 0.3% Triton (PBSTX), all with gentle shaking. Retinas were then transferred onto slides using a wide bore plastic Pasteur pipette, excess PBSTX removed with absorbent tissue, and mounted using Prolong mounting media and set overnight. Retinas were imaged using a Zeiss LSM900 confocal microscope or a Keyence epifluorescence microscope.

### Tissue processing and immunohistochemistry

Mice were injected intraperitoneally with Sulfo-NHS-biotin (200 μl of 20 mg/ml Sulfo-NHS-biotin in PBS) and sacrificed after 30 min. Tissue were harvested in 4% PFA in PBS overnight and hydrated in 30% sucrose in PBS at least for 24 hours before embedding in OCT. Tissue sections of 100 μm thickness were cut using a vibratome. Tissues and retinas were stained with Cy3-Streptavidin and imaged using confocal or epifluorescence microscopes, quantified and processed with ImageJ and Adobe Photoshop.

### Immunofluorescence analysis

For cerebellum histological analysis, tissues were processed the same as above in OCT. Tissue sections of 10 μM were blocked in 5% Normal Goat Serum (Jackson ImmunoResearch, West Grove, PA) in PBS + 1%BSA + 0.3%Triton X-100 for 1 hour at room temperature. Samples were incubated at 4°C with the following primary antibodies in PBS + 1%BDS + 0.3% Triton X-100 + 0.1% NaN: hamster anti-CD31 (1:100, cat.#MAB1398Z, Millipore, Billerica, MA), rat anti-mouse PDGFRB (1:50, cat.#14-1402-82, Clone APB5, eBiosciences, San Diego, CA), rabbit anti-CLDN5 (1:100, cat.#34-1600, Thermo Fisher Scientific, MA), Rat anti-mouse PLVAP antibody, clone MECA-32 (Bio-Rad, Hercules, CA). Excess antibody was removed by rinsing 4x in PBS + 0.1% Triton X-100 for 10 min. Samples were then incubated at room temperature for 1 h with the following secondary fluorescently labeled antibodies: FITC or Cy3 goat anti-hamster IgG, FITC or Cy3 goat anti-rat IgG, FITC or Cy3 goat anti-rabbit IgG, Cy3 goat anti-mouse IgG, FITC or Cy3 streptavidin (Jackson ImmunoResearch, West Grove, PA) diluted 1:400 in PBS + 1% BSA + 0.3% Triton X-100 for 1 h at RT. Excess antibody was removed by rinsing 4x in PBS + 0.1% Triton X-100 for 10 min. Slides were mounted in Vectashield mounting medium with DAPI (Vector labs, Burlingame, CA) and imaged with an epifluorescence microscope to obtain 10x, 20x or 40x images. Immunofluorescence signal area or density was quantified by ImageJ and normalized by vessel area (CD31 signal area). Pericyte coverage was quantified by measuring the IF staining signal length of pericyte or endothelial profile with the NeuronJ plugin for ImageJ.

### Pharmacokinetics

The pharmacokinetic profiles of L6-F4-2 were assessed via intravenous (i.v.) and intraperitoneal (i.p.) routes of drug administration in adult mice (n = 3/group). Blood samples were obtained at 10 minutes, 6 hours, 24 hours, and days 2, 4, 7, and 14 after intravenous injection of L6-F4-2 at 3 mg/kg. Similarly, blood samples were obtained at 2 hours, 6 hours, 24 hours, and days 2, 4, 7, and 14 after intraperitoneal injection of L6-F4-2 at 3 mg/kg. Serum was obtained from blood samples through centrifugation in MiniCollect® Tube 0.8 ml Z Serum Separator tubes (Greiner Bio-One). Serum samples were then assessed for L6-F4-2 using a human IgG ELISA Kit (ab195215, Abcam, MA), per manufacturer’s protocol. Pharmacokinetic parameters were estimated using noncompartmental analysis of serum L6-F4-2 concentrations in individual mice with sparse sampling using Phoenix WinNonlin. All animal experiments were performed according to national ethical guidelines in addition to the guidance and approval by the Institutional Animal Care and Use Committee (IACUC) of Surrozen, Inc. Euthanasia was conducted in compliance with the current requirements of The Guide for the Care and Use of Laboratory Animals, 8^th^ Edition, and AVMA Guidelines on Euthanasia. All animals were obtained from the Jackson Laboratory (Bar Harbor, ME). On arrival, animals were randomly assigned to group (5 animals/cage) housing and provided with rodent diet and water ad libitum. All mice were maintained on a 12:12-h light/dark photoperiod at an ambient temperature of 22 ± 2 °C.

### FACS sorting of retina vessel ECs

Retinas of P8 mice were harvested and pooled as described above. Fresh retinas were minced and incubated in 5 ml DMEM containing 200 U/ml collagenase I (Invitrogen) for 45 mins at 37°C with occasional shaking followed by filtering through a 40 μm nylon mesh. The cells were then centrifuged at 94 × g for 5 mins at 4°C and resuspended in PBS with 0.1% BSA + 2 mM EDTA. Endothelial cells were labeled with PE-Cy7 rat anti-mouse CD31 (#25-0311-82, eBiosciences, CA). PE rat anti-mouse CD45 (#516087, BD Pharmingen, CA) and 7-AAD (Invitrogen, Waltham, MA) were added to exclude hematopoietic lineage cells and dead cells. Staining was performed for 1 h at 4°C. Retina ECs were sorted into DMEM medium with an Aria II sorter (BD) at the Stanford University Shared FACS Facility and FACS data were analyzed using FlowJo software (TreeStar). Cells were further processed with single cell RNA seq Kit (10X Genomics, Pleasanton, CA) or with the Arcturus PicoPure RNA Isolation Kit (Applied Biosystems) for RNA extraction.

### Transient middle cerebral artery occlusion (tMCAO)

Adult WT C57Bl/6J mice (males, 6-8 weeks old, JAX) underwent transient middle cerebral artery occlusion by the intraluminal suture method for 45 min (body temperature maintained at 37 ±0.5 °C), as previously described^67^. Animals had free access to food and water throughout the reperfusion period. Neurological score was evaluated at 48-h post-tMCAO by a blinded observer: 0, normal motor function; 1, flexion of torso and the contralateral forelimb upon lifting by the tail; 2, circling to the ipsilateral side but normal posture at rest; 3, leaning to the ipsilateral side at rest; 4, unable to walk spontaneously. After neurological testing, the mice were deeply anesthetized with isoflurane and the brains were sectioned into 2-mm-thick sections and placed in 2% 2,3,5 triphenyltetrazolium chloride (TTC, cat. #T8877, Sigma-Aldrich, St. Louis, MO) for 10 min at 37 °C to delineate infarcts. Infarct areas were determined using an image analysis system (ImageJ). Edema was calculated by the RICH technique by assessing fractional increase in volume of the stroke vs. ipsilateral non-stroke hemispheres^68^. The persons performing the surgery and recording neurological scores were blinded to treatment status.

### Bulk RNA sequencing

For retina ECs of WT and *Ndp^KO^* mice with PBS or L6-F4-2 treatment, total RNA was extracted using the Arcturus PicoPure RNA Isolation Kit (Applied Biosystems). RNA-seq libraries were generated with the NEBNext Ultra II Directional RNA Library Prep Kit coupled with Poly(A) mRNA Magnetic Isolation Module and NEBNext multiplex oligos for Illumina (New England Biolabs). Deep sequencing was performed on the NextSeq 500 sequencing system (Illumina) with 75-cycle, paired-end sequencing. RNA data were aligned to Ensemble mouse genome using Kallisto (v 0.44.0) with default parameters. Change in gene expression between two conditions was defined as significant if |log_2_FC|>0.5 and adjusted *P* value <0.05. Complex-Heatmap was used to produce heat maps^69^.

### Single cell RNA sequencing (scRNA-seq)

WT and *Ndp^KO^* mice were treated with PBS or L6-F4-2 at P0. Retinas were harvested and retinal ECs were FACS sorted in DMEM without EDTA at P8. scRNA-seq with the 10X Genomics Single Cell 3’ platform was performed per manufacturer’s protocol. Cell capture, library preparation, and sequencing were performed as described^70^.

### Transcriptional regulatory network analysis

A transcriptional regulatory network was reverse engineered using the WT data. Metacells were generated by summing the raw counts of each with those of its five nearest neighbors in SCT-normalized gene expression, using √1-*ρ* as the distance metric, where *ρ* is the spearman correlation between samples. The data was then randomly subset to 500 such metacells, and the data was CPM normalized.

This CPM-normalized matrix of metacells was then used as input to ARACNe-AP^51^. Four regulator sets were used: TFs, coTFs, signaling, and surface proteins. ARACNe-AP was run using a ρ-value threshold of 1E-8 with 200 bootstraps for each set of regulators, with the final network being consolidated from each of the four regulator-specific networks. The final network contained 3,145 regulators with at least 50 edges that could be used for VIPER^52^ analysis.

The anchored data object consisting of joined data from the WT, *Ndp^KO^*, and *Ndp^KO^* +L6-F4-2 samples was used as input to VIPER within the PISCES pipeline. The scaled data was extracted and used as a gene expression signature for input to VIPER along with the previously generated network. The final VIPER matrix consisted of protein activity inference for 955 proteins across the 4,315 cells in the anchored dataset.

### Gene Expression Analysis

scRNA-seq data from each of the wild type with PBS treatment, *Ndp^KO^* with PBS treatment, and *Ndp^KO^* with L6-F4-2 treatment was quality controlled based on the minimum and maximum depth, minimum gene count, and maximum mitochondrial percentage values given in Supplementary Table 2. QC plots are displayed in Supplementary Fig. 5. Filtered counts were then loaded into Seurat objects following their standard pipeline. Data from the three experimental conditions were integrated with the Seurat anchoring pipeline. A UMAP reduction and clustering solution were generated with default parameters. Differential markers were identified with the ‘find_markers’ function for each cluster.

### Gene Expression Visualization

We visualized the expression of genes from two different sources in the literature^49^. First, we visualized the expression of the 13 genes they used to initially identify sub-types in the endothelial cells. We then also visualized the top five differentially expressed markers for each cell type from their data. Because the previous described analysis^49^ was performed at both P6 and P10 and our data is from P8, we took the intersection of their differentially expressed genes (see Supplementary Tables S3A and S3B in ref. ^49^ for the entire gene sets). Heatmaps were generated using the ‘complexHeatmap’ package in R, and all other visualizations were generated using ‘ggplot2’.

### NaRnEA Analysis

To assist in mapping clusters to cell types, we utilized the same gene sets from Zarkada et. al. (see Supplementary Table S4A in ref. ^49^) to perform enrichment analysis using the NaRnEA algorithm^50^. We parametrized our regulon by setting the association weight (AW) and association mode (AM) to one for all genes in the given sets. A signature was generated for each cluster via gene-level Wilcoxon Rank-Sum test for each cluster versus all others. Gene expression signature (GES) normalized enrichment scores (NES) were generated by transforming the p-value to a Z-score using ‘qnorm’, then signing this Z-score based on the rank-biserial correlation. NaRnEA was then run to identify enrichment of each gene set in the signature for each cluster.

### Statistical analysis of cell type frequency by experimental condition

A chi-square test was performed between the WT and *Ndp^KO^* cell-type vectors as well as between the *Ndp^KO^* and *Ndp^KO^* +L6-F4-2 cell-type vectors using the ‘chisq.test’ function in R. To extract cell-type level conclusions, we analyzed the standardized residuals for each group. P-values were corrected with the Benjamini-Hochberg procedure.

### Statistical analysis

Statistical analysis was performed using GraphPad Prism. All statistical tests used biological replicates and are indicated by group size (n) in figure legends. Results were expressed as mean ± s.e.m. (standard error of the mean). *p<0.05, **p<0.01, Mann-Whitney U test.

---

**Supplementary Figure 1.**
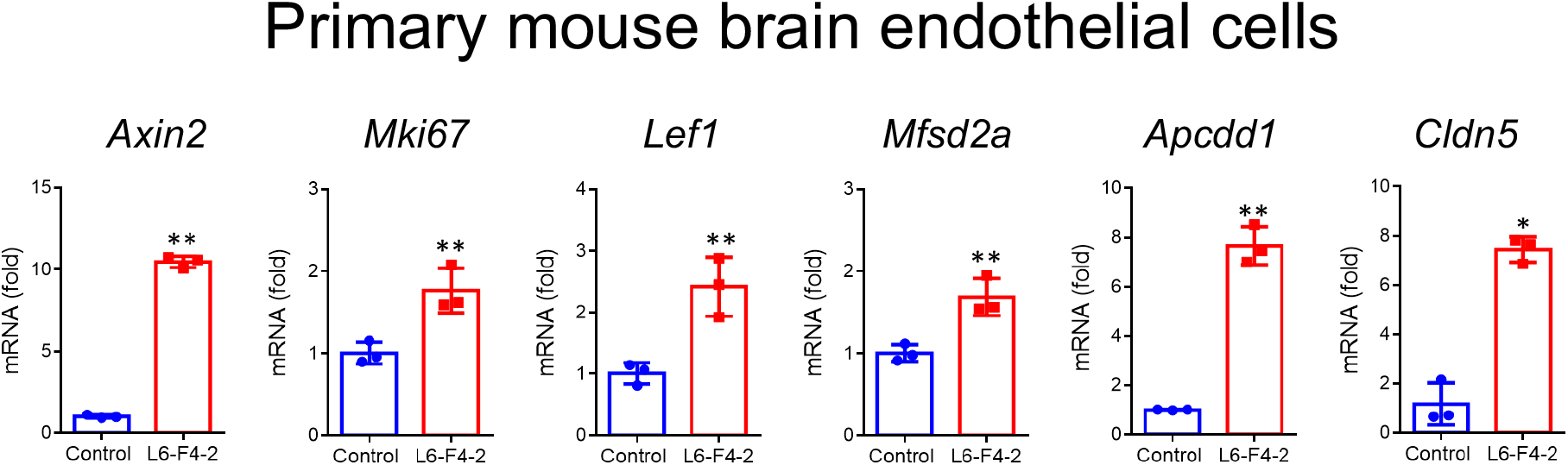
L6-F4-2-stimulated gene expression in primary mouse brain endothelial cells. Primary brain endothelial cells were isolated from adult C57Bl/6J mice and treated with NIST or L6-F4-2 (10 nM) for 24 hours. Cells were harvested and expression of indicated genes were quantified by qRT-PCR. Error bars represent mean ± s.e.m., n=3, *p < 0.05, **p<0.01, Mann-Whitney U test.

**Supplementary Figure 2.**
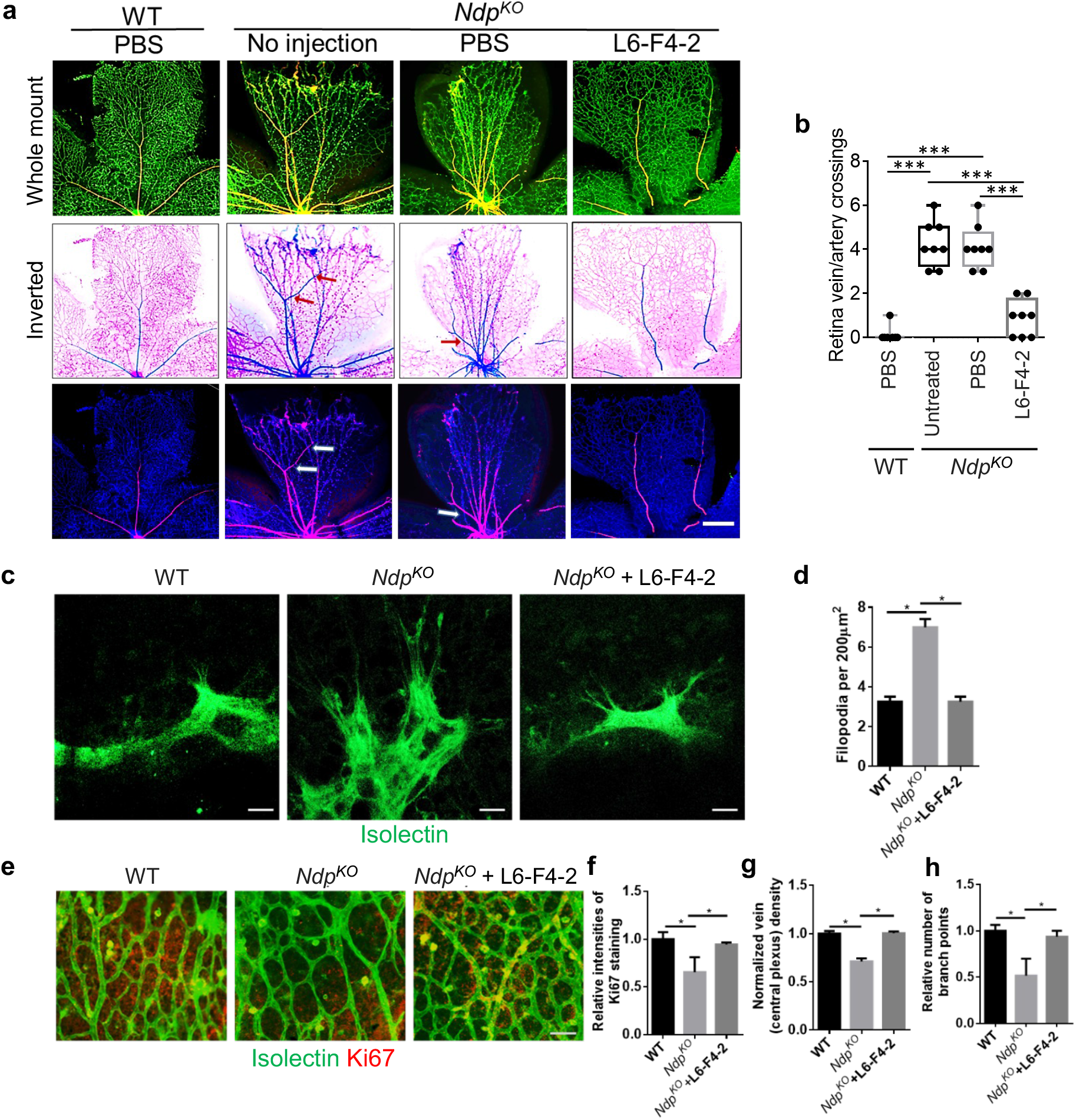
Rescue of *Ndp^KO^* retinal vascular phenotypes by L6-F4-2. WT (control) and *Ndp^KO^* mice were treated with PBS or L6-F4-2 by intravitreal injection (0.19 μg) at P0 and retinas were harvested at P8. (**a** and **b**) Isolectin B4 labeled veins (green/blue) and smooth muscle labeled arteries (yellow/red) at P8 retina and quantification of crossings. Aberrant artery/vein crossings are seen in *Ndp^KO^* and PBS treated *Ndp^KO^* retinas (arrows), but not in the L6-F4-2-treated group. Error bars represent mean ± s.e.m., *Ndp* WT n=7, *Ndp^KO^* n=8, ***p < 0.001; Mann-Whitney U test. **c** P8 retinas were stained with isolectin and filopodia quantified (**d**) in WT, *Ndp^KO^* and *Ndp^KO^+L6-* F4-2 mice. Scale bar 200 μm. **e** Retina whole mounts were stained with isolectin and anti-Ki67 and the relative intensities of Ki67 staining are quantified in (**f**). Quantification of vein density (central plexus) (**g**) and branch point numbers (**h**). Scale bar 100 μm. Error bars represent mean ± s.e.m., n=3, *p<0.05, Mann-Whitney U test.

**Supplementary Figure 3.**
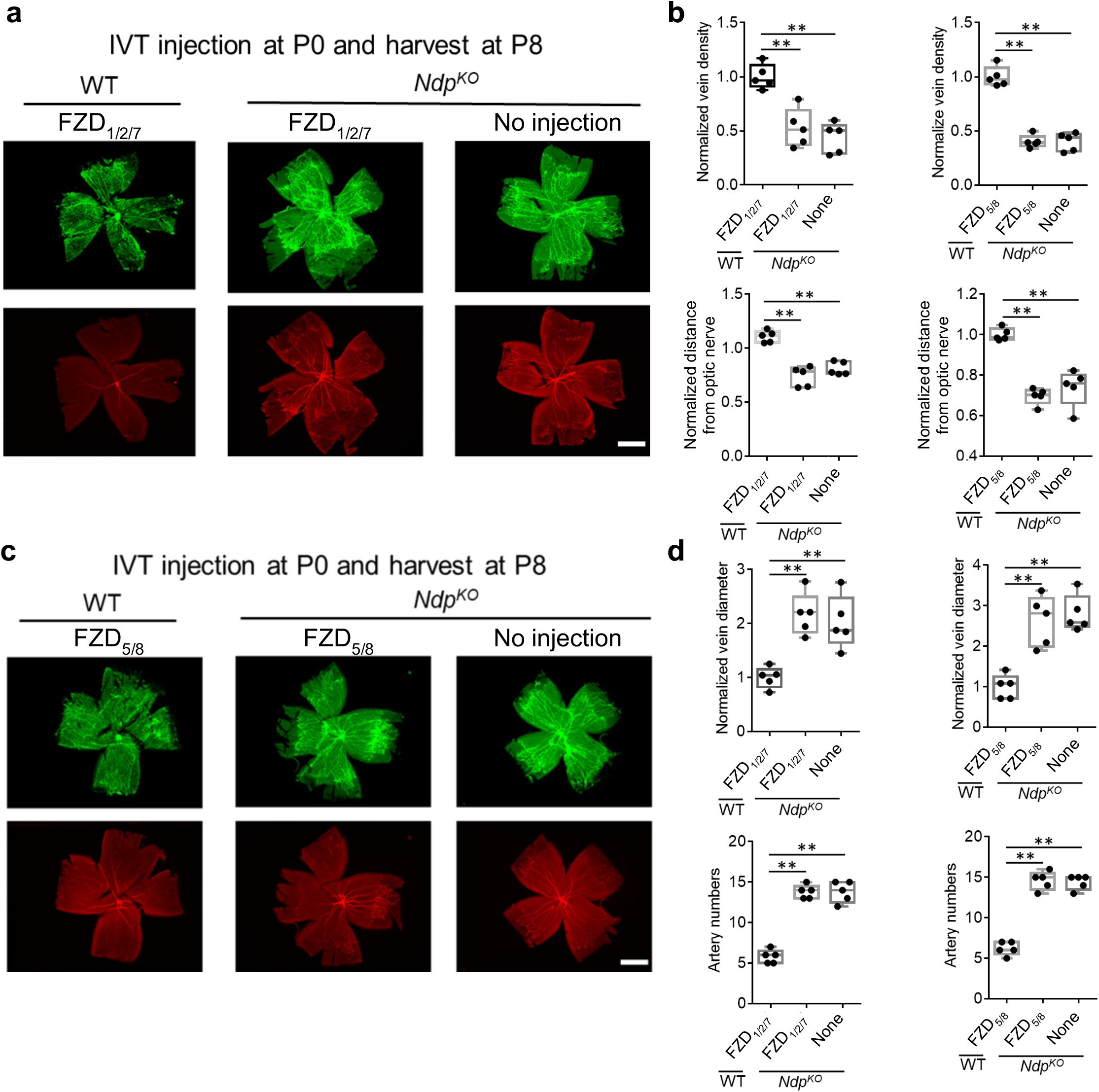
FZD1/2/7-and FZD5/8-selective surrogates did not rescue retinal developmental angiogenesis in *Ndp^KO^* mice. WT (control) and *Ndp^KO^* mice were treated with PBS or FZD surrogates by intravitreal (IVT) injection (0.19 μg) at P0 with retina harvest at P8. FZD1/2/7 surrogate (**a and b)** and FZD5/8 surrogate (**c and d**) did not rescue abnormal retinal architectures. Isolectin B4-labeled veins (green) and anti-smooth muscle actin-labeled arteries (red) are depicted in retinal flat mounts. Scale bar = 0.5 mm. Error bars represent mean ± s.e.m., **p<0.01, Mann-Whitney U test.

**Supplementary Figure 4.**
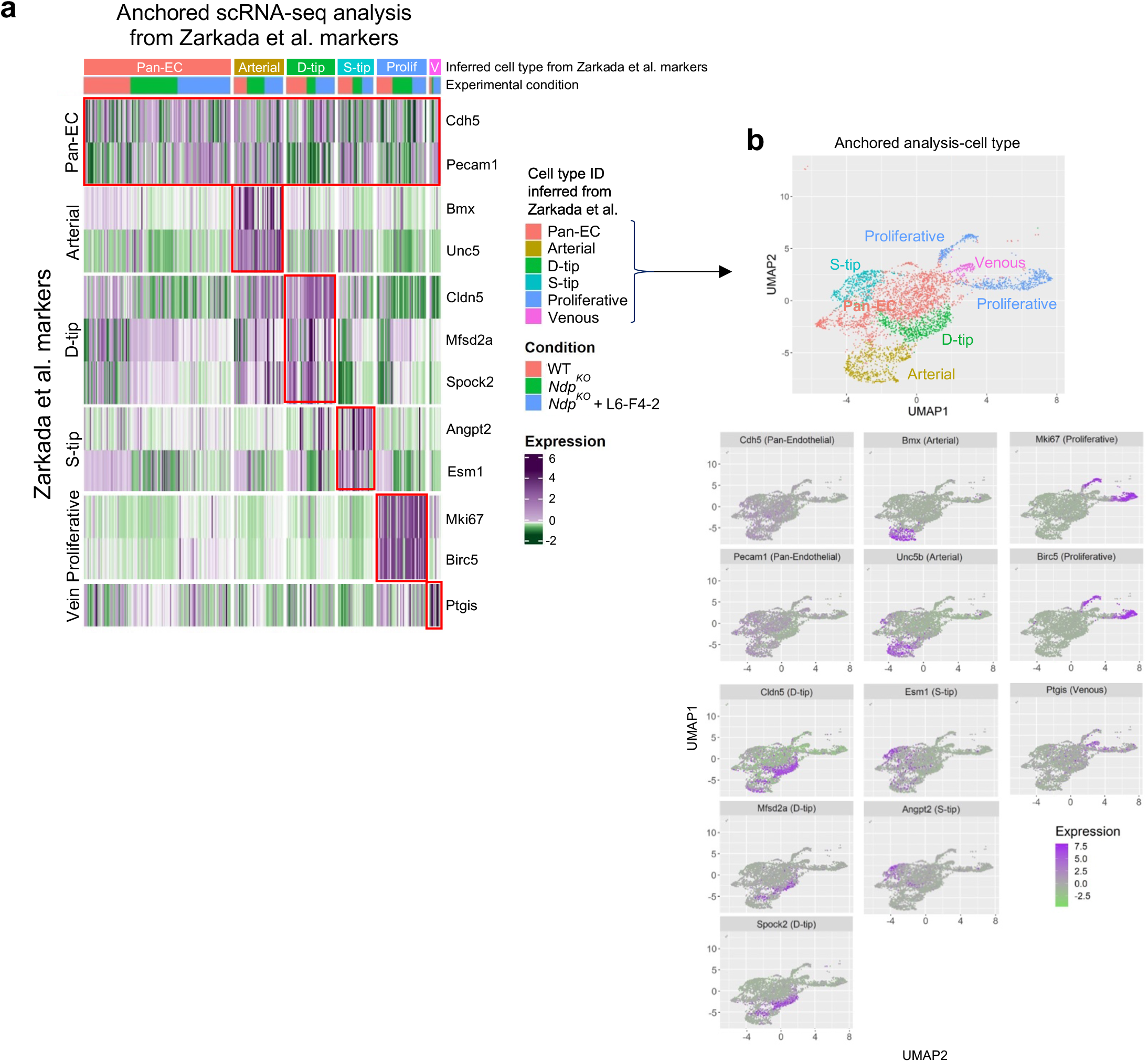
**a** Heat map analysis of scRNA-seq data from Fig. 3d, displaying for each cluster SCT-normalized expression of cell type markers from Zarkada et al. (ref. ^49^) in a Seurat-integrated dataset, containing all three of the WT, *Ndp^KO^*, and *Ndp^KO^* + L6-F4-2 experimental conditions. Concordance between Zarkada et al. markers and our scRNA-seq Seurat clusters identified *Cldn5^+^Mfsd2a^+^Spock2^+^* D-tip, *Angpt2^+^Esm1^+^* S-tip, *Unc5b^+^Bmx^+^* arterial, *Ptgis^high^* venous and *Mki67^+^Birc5^+^* proliferative endothelium in our scRNA-seq dataset. The remaining cells expressing *Cdh5* and *Pecam1* were described as pan-endothelial (pan-EC). **b** Feature plots labeling expression of the indicated genes within the pan-EC, D-tip, S-tip, arterial, proliferative, and venous populations identified from (**a**) in UMAP space.

**Supplementary Figure 5.**
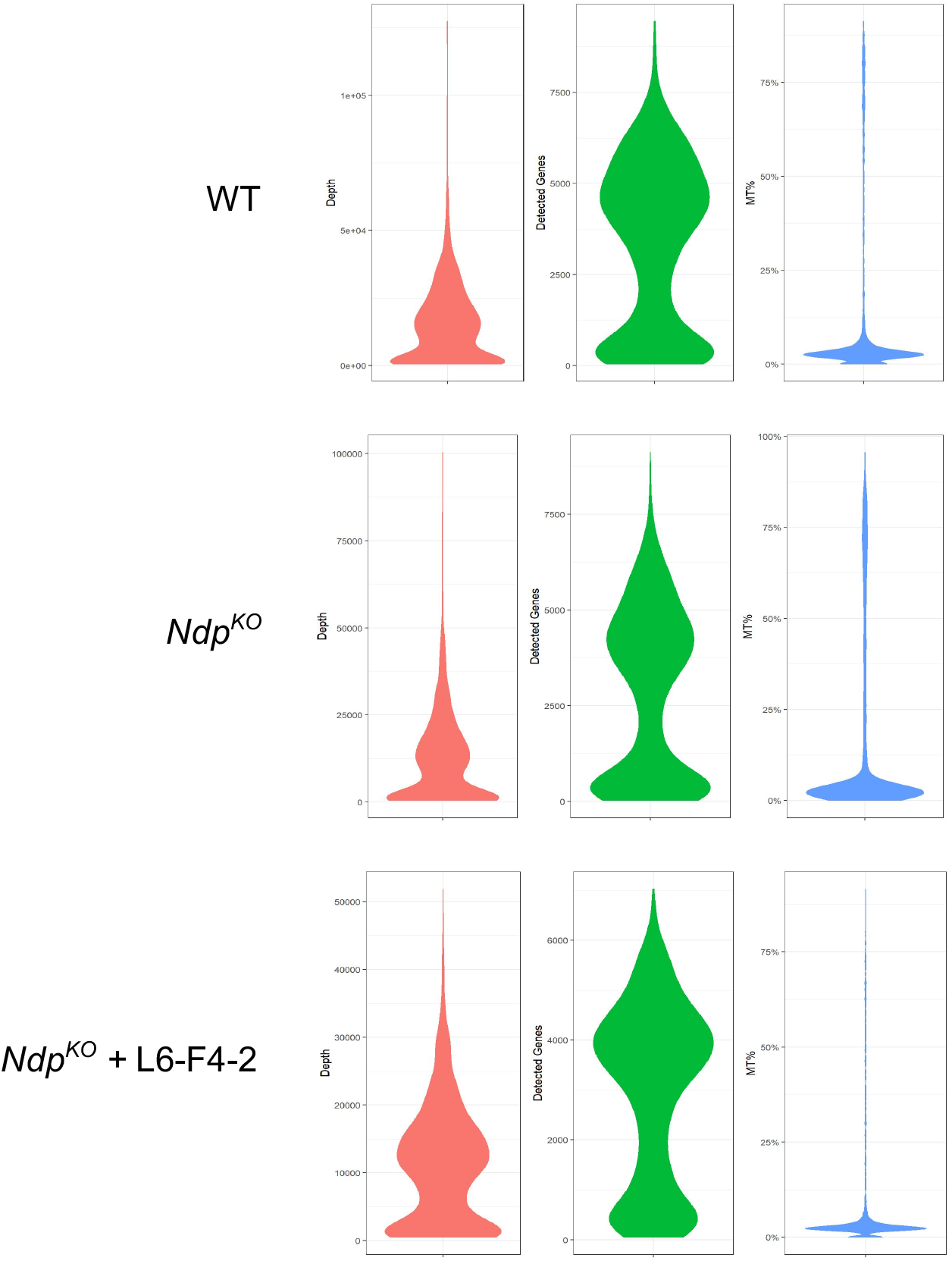
Quality control metrics for scRNA-seq for each of the three experimental conditions. Values selected for filtration were selected based on this figure and can be found in Supplementary Table 1.

**Supplementary Figure 6.**
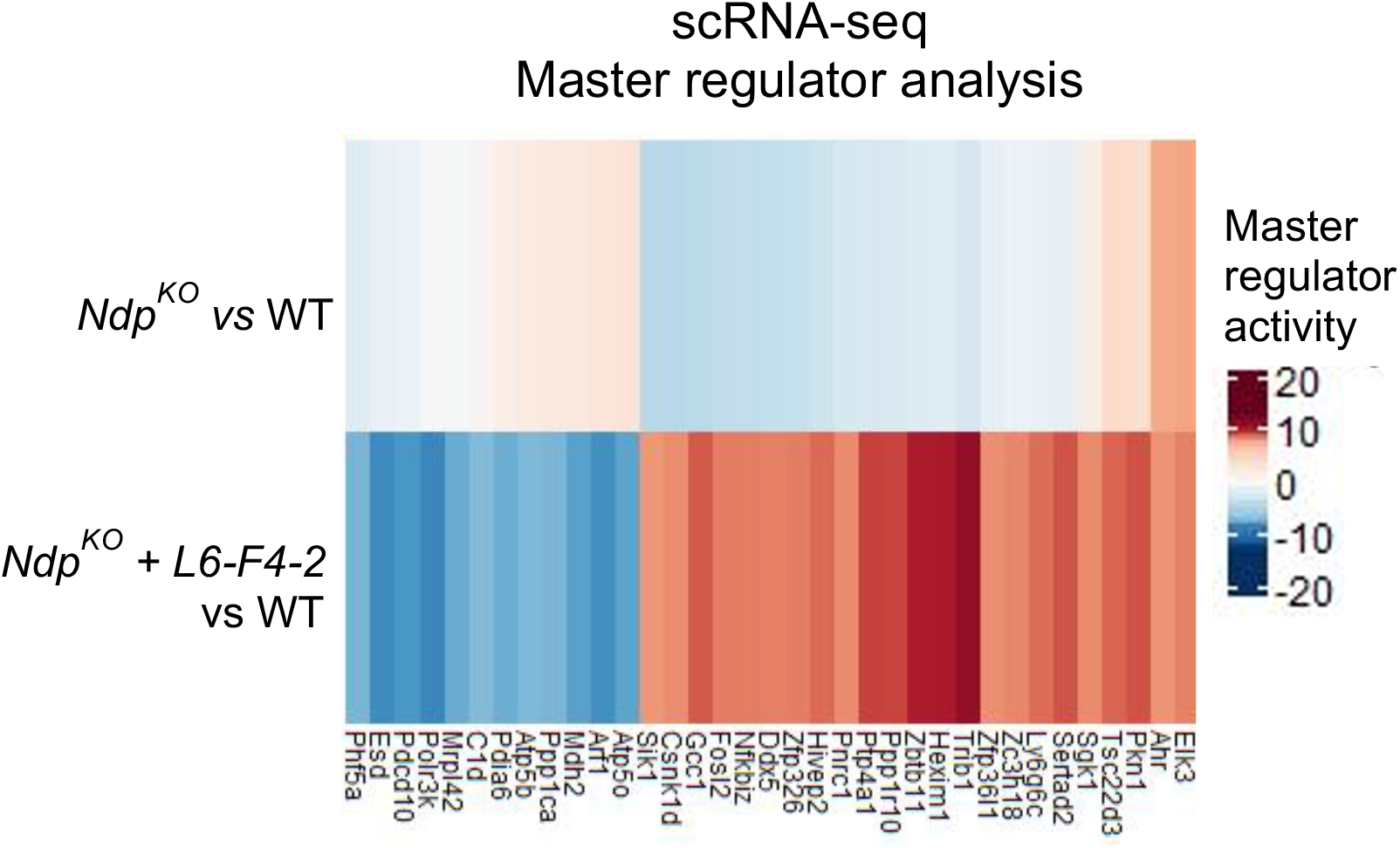
Master regulators of the *Ndp^KO^* + L6-F4-2 condition versus WT as identified by PISCES analysis of scRNA-seq data without significant activity in the *Ndp^KO^* condition. These proteins represent transcriptional programs driven solely or in large part by the therapeutic agent rather than the Norrin knockout phenotype.

**Supplementary Figure 7.**
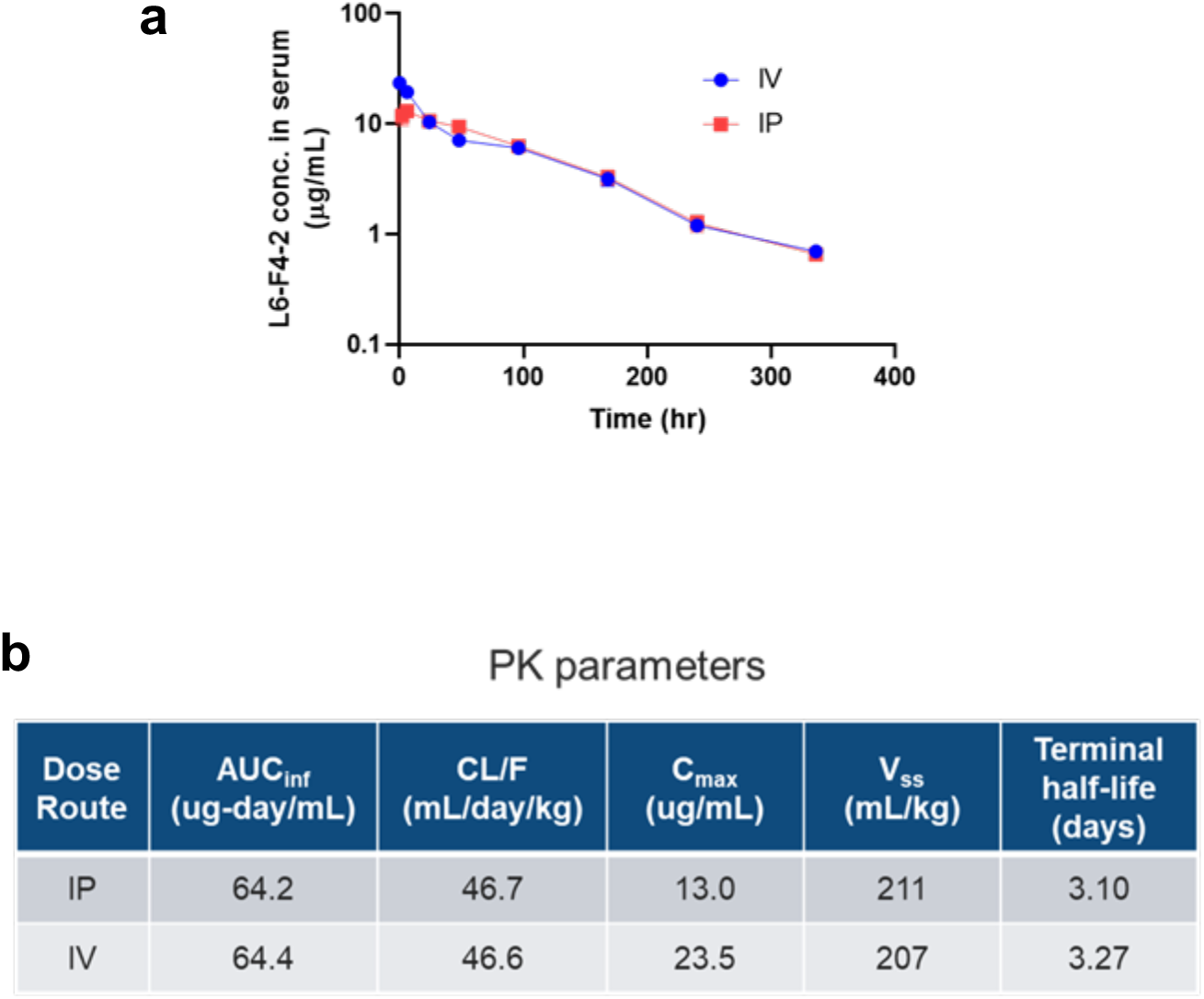
Pharmacokinetics of L6-F4-2. **a** ELISA determination of the concentration-time profile of L6-F4-2 in wild-type mouse serum following a single 3 mg/kg intravenous (IV) or intraperitoneal (IP) dose (Mean ± SEM, n=3 per time point). **b** Mean PK parameters of L6-F4-2. F = bioavailability, AUC_inf_ = area under the curve to time infinity, CL = clearance, V_ss_ = volume of distribution at steady state, C_max_ = maximum observed concentration. The bioavailability after IP dosing was 100%.

**Supplementary Figure 8.**
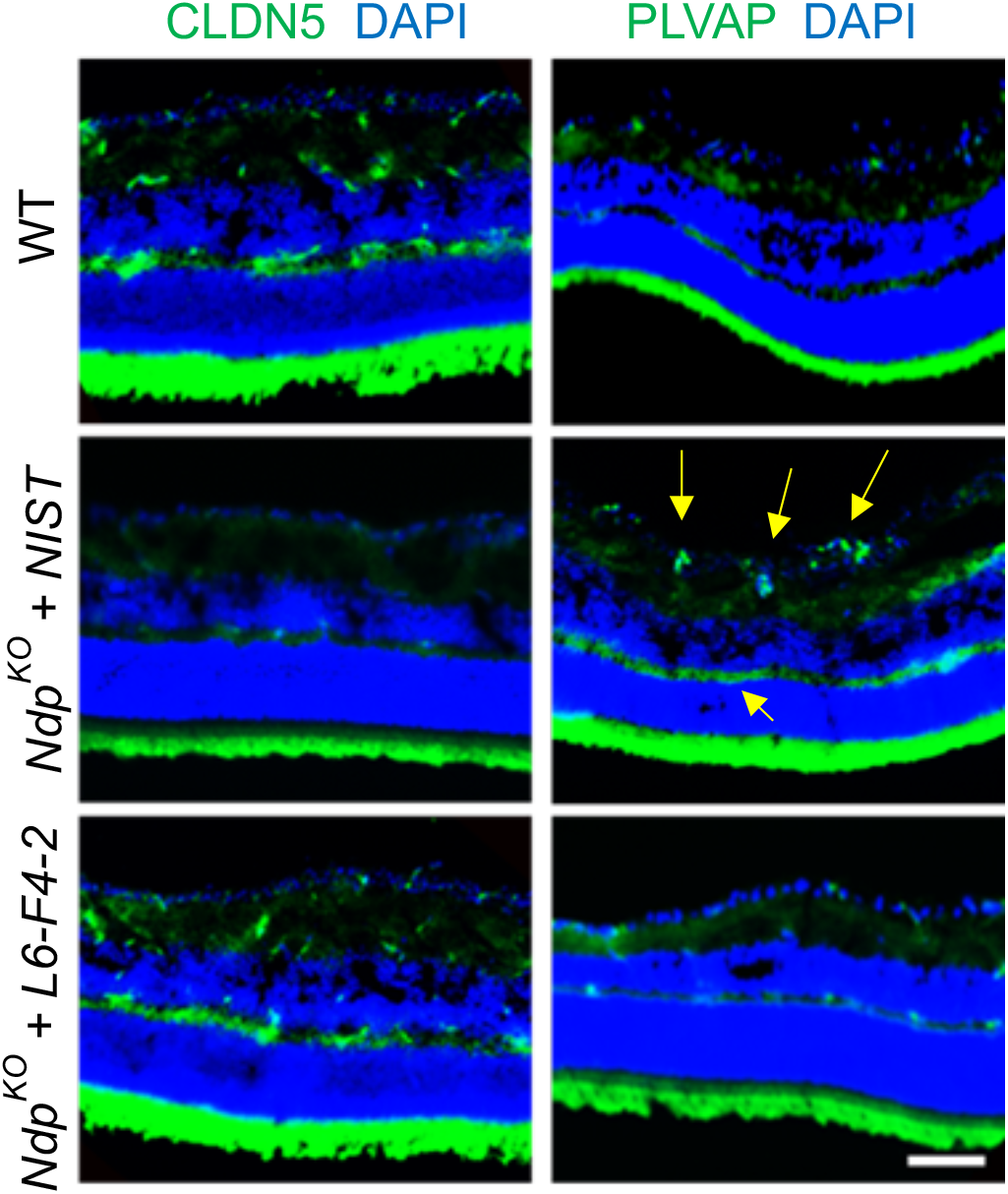
L6-F4-2 administration increased the expression of CLDN5 and decreased expression of PLVAP in *Ndp^KO^* mice. *Ndp*^KO^ mice were treated with NIST or L6-F4-2 (2.5 mg/kg, i.p.) at P0, P7, P14 and retinas were harvested at P21. *Ndp^KO^* mice lacked endothelial CLDN5 in the superficial and intermediate retinal layers, which was rescued by L6-F4-2. In contrast, *Ndp*^KO^ mice inappropriately expressed PLVAP in the superficial and intermediate retinal layer (yellow arrows), which was rescued by L6-F4-2. Scale bar 100 μm.

**Supplementary Table 1.** Filtration values used to subset scRNA-seq raw data in Seurat, based on QC metrics visualized in Supplementary Figure 5.

|  | WT | *Ndp*<sup>KO</sup> | *Ndp*<sup>KO</sup> +L6-F4-2 |
| --- | --- | --- | --- |
| **nCount_RNA** | >1250, <75000 | >1000 | >3000 |
| **nFeature_RNA** | >1000 | >1250 | >1000 |
| **Mt.features** | <12.5% | <25% | <12.5% |

**Supplementary Table 2.** QC filtering values for each scRNA-seq experimental condition.

|  | min.depth | max.depth | min.genes | max.mat |
| --- | --- | --- | --- | --- |
| WT | 2000 | 50000 | 1000 | 0.1 |
| *Ndp*<sup>KO</sup> | 2000 | 50000 | 1000 | 0.1 |
| *Ndp*<sup>KO</sup> + L6-F4-2 | 2000 | 40000 | 1000 | 0.1 |

## REFERENCES

1. Claesson-Welsh, L., Dejana, E. & McDonald, D.M. Permeability of the Endothelial Barrier: Identifying and Reconciling Controversies. Trends Mol Med 27, 314–331 (2021).

2. Horng, S., et al. Astrocytic tight junctions control inflammatory CNS lesion pathogenesis. J Clin Invest 127, 3136–3151 (2017).

3. Rosenberg, G.A. Neurological diseases in relation to the blood-brain barrier. J Cereb Blood Flow Metab (2012).

4. Buzhdygan, T.P., et al. The SARS-CoV-2 spike protein alters barrier function in 2D static and 3D microfluidic in vitro models of the human blood-brain barrier. bioRxiv (2020).

5. Larochelle, C., Alvarez, J.I. & Prat, A. How do immune cells overcome the blood-brain barrier in multiple sclerosis? FEBS Lett 585, 3770–3780 (2011).

6. Zenaro, E., Piacentino, G. & Constantin, G. The blood-brain barrier in Alzheimer’s disease. Neurobiol Dis 107, 41–56 (2017).

7. Zhao, Z., Nelson, A.R., Betsholtz, C. & Zlokovic, B.V. Establishment and Dysfunction of the Blood-Brain Barrier. Cell 163, 1064–1078 (2015).

8. Snellings, D.A., et al. Cerebral Cavernous Malformation: From Mechanism to Therapy. Circ Res 129, 195–215 (2021).

9. Schuijers, J., Mokry, M., Hatzis, P., Cuppen, E. & Clevers, H. Wnt-induced transcriptional activation is exclusively mediated by TCF/LEF. EMBO J 33, 146–156 (2014).

10. Wang, Y., et al. Interplay of the Norrin and Wnt7a/Wnt7b signaling systems in blood-brain barrier and blood-retina barrier development and maintenance. Proc Natl Acad Sci U S A 115, E11827–E11836 (2018).

11. Zhou, Y., et al. Canonical WNT signaling components in vascular development and barrier formation. J Clin Invest 124, 3825–3846 (2014).

12. Daneman, R., et al. Wnt/beta-catenin signaling is required for CNS, but not non-CNS, angiogenesis. Proc Natl Acad Sci U S A 106, 641–646 (2009).

13. Liebner, S., et al. Wnt/beta-catenin signaling controls development of the blood-brain barrier. J Cell Biol 183, 409–417 (2008).

14. Eubelen, M., et al. A molecular mechanism for Wnt ligand-specific signaling. Science 361(2018).

15. Chang, J., et al. Gpr124 is essential for blood-brain barrier integrity in central nervous system disease. Nat Med 23, 450–460 (2017).

16. Guerit, S., et al. Astrocyte-derived Wnt growth factors are required for endothelial blood-brain barrier maintenance. Prog Neurobiol 199, 101937 (2021).

17. Corada, M., et al. Fine-Tuning of Sox17 and Canonical Wnt Coordinates the Permeability Properties of the Blood-Brain Barrier. Circ Res 124, 511–525 (2019).

18. Wang, Y., et al. Norrin/Frizzled4 signaling in retinal vascular development and blood brain barrier plasticity. Cell 151, 1332–1344 (2012).

19. Zhang, C., et al. Norrin-induced Frizzled4 endocytosis and endo-lysosomal trafficking control retinal angiogenesis and barrier function. Nat Commun 8, 16050 (2017).

20. Chang, T.H., et al. Structure and functional properties of Norrin mimic Wnt for signalling with Frizzled4, Lrp5/6, and proteoglycan. Elife 4(2015).

21. Xu, Q., et al. Vascular development in the retina and inner ear: control by Norrin and Frizzled-4, a high-affinity ligand-receptor pair. Cell 116, 883–895 (2004).

22. Richter, M., et al. Retinal vasculature changes in Norrie disease mice. Invest Ophthalmol Vis Sci 39, 2450–2457 (1998).

23. Rehm, H.L., et al. Vascular defects and sensorineural deafness in a mouse model of Norrie disease. J Neurosci 22, 4286–4292 (2002).

24. Ye, X., et al. Norrin, frizzled-4, and Lrp5 signaling in endothelial cells controls a genetic program for retinal vascularization. Cell 139, 285–298 (2009).

25. Xia, C.H., Yablonka-Reuveni, Z. & Gong, X. LRP5 is required for vascular development in deeper layers of the retina. PLoS One 5, e11676 (2010).

26. Chen, J., et al. Retinal expression of Wnt-pathway mediated genes in low-density lipoprotein receptor-related protein 5 (Lrp5) knockout mice. PLoS One 7, e30203 (2012).

27. Zhang, C., et al. Endothelial Cell-Specific Inactivation of TSPAN12 (Tetraspanin 12) Reveals Pathological Consequences of Barrier Defects in an Otherwise Intact Vasculature. Arterioscler Thromb Vasc Biol 38, 2691–2705 (2018).

28. Chidiac, R., et al. A Norrin/Wnt surrogate antibody stimulates endothelial cell barrier function and rescues retinopathy. EMBO Mol Med 13, e13977 (2021).

29. Ye, X., Wang, Y. & Nathans, J. The Norrin/Frizzled4 signaling pathway in retinal vascular development and disease. Trends in molecular medicine 16, 417–425 (2010).

30. Ohlmann, A. & Tamm, E.R. Norrin: molecular and functional properties of an angiogenic and neuroprotective growth factor. Prog Retin Eye Res 31, 243-257 (2012).

31. Perez-Vilar, J. & Hill, R.L. Norrie disease protein (norrin) forms disulfide-linked oligomers associated with the extracellular matrix. J Biol Chem 272, 33410–33415 (1997).

32. Niehrs, C. Norrin and frizzled; a new vein for the eye. Dev Cell 6, 453–454 (2004).

33. Gao, X. & Hannoush, R.N. Single-cell imaging of Wnt palmitoylation by the acyltransferase porcupine. Nature chemical biology 10, 61–68 (2014).

34. Janda, C.Y. & Garcia, K.C. Wnt acylation and its functional implication in Wnt signalling regulation. Biochemical Society transactions 43, 211–216 (2015).

35. Yan, K.S., et al. Non-equivalence of Wnt and R-spondin ligands during Lgr5(+) intestinal stem-cell self-renewal. Nature 545, 238–242 (2017).

36. Janda, C.Y., et al. Surrogate Wnt agonists that phenocopy canonical Wnt and beta-catenin signalling. Nature 545, 234–237 (2017).

37. Dang, L.T., et al. Receptor subtype discrimination using extensive shape complementary designed interfaces. Nat Struct Mol Biol 26, 407–414 (2019).

38. Miao, Y., et al. Next-Generation Surrogate Wnts Support Organoid Growth and Deconvolute Frizzled Pleiotropy In Vivo. Cell Stem Cell 27, 840–851 e846 (2020).

39. Tao, Y., et al. Tailored tetravalent antibodies potently and specifically activate Wnt/Frizzled pathways in cells, organoids and mice. Elife 8(2019).

40. Chen, H., et al. Development of Potent, Selective Surrogate WNT Molecules and Their Application in Defining Frizzled Requirements. Cell chemical biology 27, 598–609 e594 (2020).

41. Ke, J., et al. Structure and function of Norrin in assembly and activation of a Frizzled 4-Lrp5/6 complex. Genes Dev 27, 2305–2319 (2013).

42. Bang, I., et al. Biophysical and functional characterization of Norrin signaling through Frizzled4. Proc Natl Acad Sci U S A 115, 8787–8792 (2018).

43. Wang, Y., et al. Beta-catenin signaling regulates barrier-specific gene expression in circumventricular organ and ocular vasculatures. Elife 8(2019).

44. Ben-Zvi, A., et al. Mfsd2a is critical for the formation and function of the blood-brain barrier. Nature 509, 507–511 (2014).

45. Wang, Z., et al. Wnt signaling activates MFSD2A to suppress vascular endothelial transcytosis and maintain blood-retinal barrier. Sci Adv 6, eaba7457 (2020).

46. Berger, W., et al. Isolation of a candidate gene for Norrie disease by positional cloning. Nat Genet 1, 199–203 (1992).

47. Zuercher, J., Fritzsche, M., Feil, S., Mohn, L. & Berger, W. Norrin stimulates cell proliferation in the superficial retinal vascular plexus and is pivotal for the recruitment of mural cells. Hum Mol Genet 21, 2619-2630 (2012).

48. Laksitorini, M.D., Yathindranath, V., Xiong, W., Hombach-Klonisch, S. & Miller, D.W. Modulation of Wnt/beta-catenin signaling promotes blood-brain barrier phenotype in cultured brain endothelial cells. Sci Rep 9, 19718 (2019).

49. Zarkada, G., et al. Specialized endothelial tip cells guide neuroretina vascularization and blood-retina-barrier formation. Dev Cell 56, 2237–2251 e2236 (2021).

50. A. Griffin, L.V. C. Chiuzan, A. Califano. An Information Theoretic Framework for Protein Activity Measurement. (bioRxiv, 2022).

51. Lachmann, A., Giorgi, F.M., Lopez, G. & Califano, A. ARACNe-AP: gene network reverse engineering through adaptive partitioning inference of mutual information. Bioinformatics 32, 2233–2235 (2016).

52. Alvarez, M.J., et al. Functional characterization of somatic mutations in cancer using network-based inference of protein activity. Nat Genet 48, 838–847 (2016).

53. Bosma, E.K., van Noorden, C.J.F., Schlingemann, R.O. & Klaassen, I. The role of plasmalemma vesicle-associated protein in pathological breakdown of blood-brain and blood-retinal barriers: potential novel therapeutic target for cerebral edema and diabetic macular edema. Fluids Barriers CNS 15, 24 (2018).

54. Gastfriend, B.D., et al. Wnt signaling mediates acquisition of blood-brain barrier properties in naive endothelium derived from human pluripotent stem cells. Elife 10(2021).

55. Benjamin, E.J., et al. Heart Disease and Stroke Statistics-2017 Update: A Report From the American Heart Association. Circulation 135, e146–e603 (2017).

56. Miller, D.J., Simpson, J.R. & Silver, B. Safety of thrombolysis in acute ischemic stroke: a review of complications, risk factors, and newer technologies. Neurohospitalist 1, 138–147 (2011).

57. Saver, J.L. Hemorrhage after thrombolytic therapy for stroke: the clinically relevant number needed to harm. Stroke 38, 2279–2283 (2007).

58. Brown, D.L., Barsan, W.G., Lisabeth, L.D., Gallery, M.E. & Morgenstern, L.B. Survey of emergency physicians about recombinant tissue plasminogen activator for acute ischemic stroke. Ann Emerg Med 46, 56–60 (2005).

59. Gautam, J. & Yao, Y. Roles of Pericytes in Stroke Pathogenesis. Cell Transplant 27, 1798–1808 (2018).

60. Cao, L., Zhou, Y., Chen, M., Li, L. & Zhang, W. Pericytes for Therapeutic Approaches to Ischemic Stroke. Front Neurosci 15, 629297 (2021).

61. Armulik, A., et al. Pericytes regulate the blood-brain barrier. Nature 468, 557–561 (2010).

62. Junge, H.J. Ligand-Selective Wnt Receptor Complexes in CNS Blood Vessels: RECK and GPR124 Plugged In. Neuron 95, 983–985 (2017).

63. Martin, M., et al. Engineered Wnt ligands enable blood-brain barrier repair in neurological disorders. Science 375, eabm4459 (2022).

64. Zhang, Z., et al. Tissue-targeted R-spondin mimetics for liver regeneration. Sci Rep 10, 13951 (2020).

65. Talks, J., et al. The use of real-world evidence for evaluating anti-vascular endothelial growth factor treatment of neovascular age-related macular degeneration. Surv Ophthalmol 64, 707–719 (2019).

66. Lo, M., et al. Effector-attenuating Substitutions That Maintain Antibody Stability and Reduce Toxicity in Mice. J Biol Chem 292, 3900–3908 (2017).

67. Maier, C.M., Hsieh, L., Crandall, T., Narasimhan, P. & Chan, P.H. Evaluating therapeutic targets for reperfusion-related brain hemorrhage. Ann Neurol 59, 929–938 (2006).

68. Boyko, M., et al. Establishment of novel technical methods for evaluating brain edema and lesion volume in stroked rats: A standardization of measurement procedures. Brain Res 1718, 12–21 (2019).

69. Gu, Z., Eils, R. & Schlesner, M. Complex heatmaps reveal patterns and correlations in multidimensional genomic data. Bioinformatics 32, 2847–2849 (2016).

70. Salahudeen, A.A., et al. Progenitor identification and SARS-CoV-2 infection in human distal lung organoids. Nature 588, 670–675 (2020).

